# Balancing Model-Based and Memory-Free Action Selection under Competitive Pressure

**DOI:** 10.1101/662700

**Authors:** Atsushi Kikumoto, Ulrich Mayr

## Abstract

In competitive situations, winning depends on selecting actions that surprise the opponent. Such unpredictable action can be generated based on representations of the opponent’s strategy and choice history (model-based counter-prediction) or by choosing actions in a memory-free, stochastic manner. Across five different experiments using a variant of a matching-pennies game with simulated and human opponents we found that people toggle between these two strategies, using model-based selection used when recent wins signal the appropriateness of the current model, but reverting to stochastic selection following losses. Also, after wins, feedback-related, mid-frontal EEG activity reflected information about the opponent’s global and local strategy, and predicted upcoming choices. After losses, this activity was nearly absent—indicating that the internal model is suppressed after negative feedback. We suggest that the mixed-strategy approach allows negotiating two conflicting goals: (1) exploiting the opponent’s deviations from randomness while (2) remaining unpredictable for the opponent.

---

Even the most powerful backhand stroke in a tennis match loses its punch when the opponent knows it is coming. Competitions that require real-time, fast-paced decision making are typically won by the player with the greatest skill in executing action plans *and* who are able to choose their moves in the least predictable manner (1–4). Yet, how people can consistently achieve the competitive edge of surprise is not well understood. The fundamental challenge towards such an unerstanding lies in the fact that our cognitive system is geared towards using memory records of the recent selection histoy to exploit regularities in the environment. However, as suggested by decades of research (5–9), these same memory records will also produce constraints on current action selection that can work against unpredictable behavior.

One such memory-based constraint on unpredictable action selection is that people often tend to repeat the last-executed action plan. A considerable body of research with the “voluntary task-switching” paradigm (8, 9) has revealed a robust perseveration bias, even when subjects are instructed to choose randomly between two different action plans on a trial-by-trial basis––a regularity that in competitions could be easily exploited by a perceptive opponent.

Another important constraint is the win-stay/lose-shift bias, that is a tendency to repeat the most recently reinforced action and abandon the most recently punished action. Reinforcement-based action selection does not require an internal representation of the task environment and is therefore often referred to as “model-free”. Previous work has revealed that reinforcement learning can explain some of the choice behavior in competitive situations (10–12). Yet, players who rely on reinforcement-based selection can also be counter-predicted by their opponent, or run the risk of missing regularities in their opponents’ behavior. Therefore, recent research indicates that when performing against sophisticated opponents, model-free choice can be replaced through model-based selection, where choices are based on a representation of task-space contingencies (13), including beliefs about the opponent’s strategies (14, 15).

Model-based selection is consistent with the view of humans as rational decision makers (2, 3), yet also has known limitations. For example, it depends on attentional and/or working memory resources that vary across and within individuals (16). In addition, people are prone to judgement and decision errors, such as the confirmation bias, that get in the way of consistently adaptive, model-based selection (17).

In light of the shortcomings of both standard, model-free choice and model-based strategies it is useful to consider the possibility that in some situations, actors can chose in a memory-free and thus stochastic manner (15). Memory-free choice would establish a “clean-slate” that prevents the system from getting stuck with a sub-optimal strategy and instead allows exploration of the full space of possible moves. Moreover, it reduces the danger of being counter-predicted by the opponent (18, 19). At the same time, an obvious drawback of stochastic choice is that without a representation of the opponent, systematic deviations from randomness in the opponent’s behavior remain undetected and therefore cannot be exploited. In addition, just as is the case for model-based selection, stochastic selection puts high demands on cognitive control resources (20) and therefore it is not clear under which circumstances people can consistently ignore or suppress context representations in order to choose in a memory-free manner (21)

As the model-based and the memory-free strategy both come with strengths and limitations, one potential solution is that people use a simple heuristic to move back and forth between these two modes of selection. Specifically, positive feedback (i.e., wins on preceding moves) could serve as a cue that the current model is adequate and should be maintained. In contrast, negative feedback might serve as a signal that the current model needs to be suspended in favor of a memory-free mode of selection that maximizes exploration and unpredictability.

In the current work, we used an experimental paradigm that provides a clear behavioral signature of model-based versus memory-free choices as a function of preceding win versus loss feedback. We found that following win feedback, people tended to choose their next move both on the basis of recent history and a more global model of the opponent. However following losses, we did not simply see choice behavior revert back towards simple memory-driven biases. Rather choices were less determined by recent history and task context––in other words more stochastic. In addition, we present neural evidence that loss feedback literally “cleans the slate” by temporarily diminishing the representation of the internal model (15, 22).

## Results

### Overview

Our experimental situation marries the voluntary task-switching paradigm with a two-person, matching-pennies game, played for real money against either simulated or human opponents. As shown in Figure 1a, on each trial players saw a circle, either on the bottom or the top of a vertically arranged rectangle. By pressing a key that is spatially compatible to the current circle location, the player could keep the circle at that location (a “freeze” choice), whereas the spatially opposite response moved the circle to the opposite location (a “run” choice). Wins versus losses were signaled through a smiley or frowny face at the place of the post-response circle position. Players were assigned either the role of the “fox” or the “rabbit”. Foxes win a given trial when they “catch the rabbit”, that is when they pick the same move as the rabbit on that trial. Rabbits win when they “escape the fox”, that is when they pick the move not chosen by the fox.

**Fig. 1.**
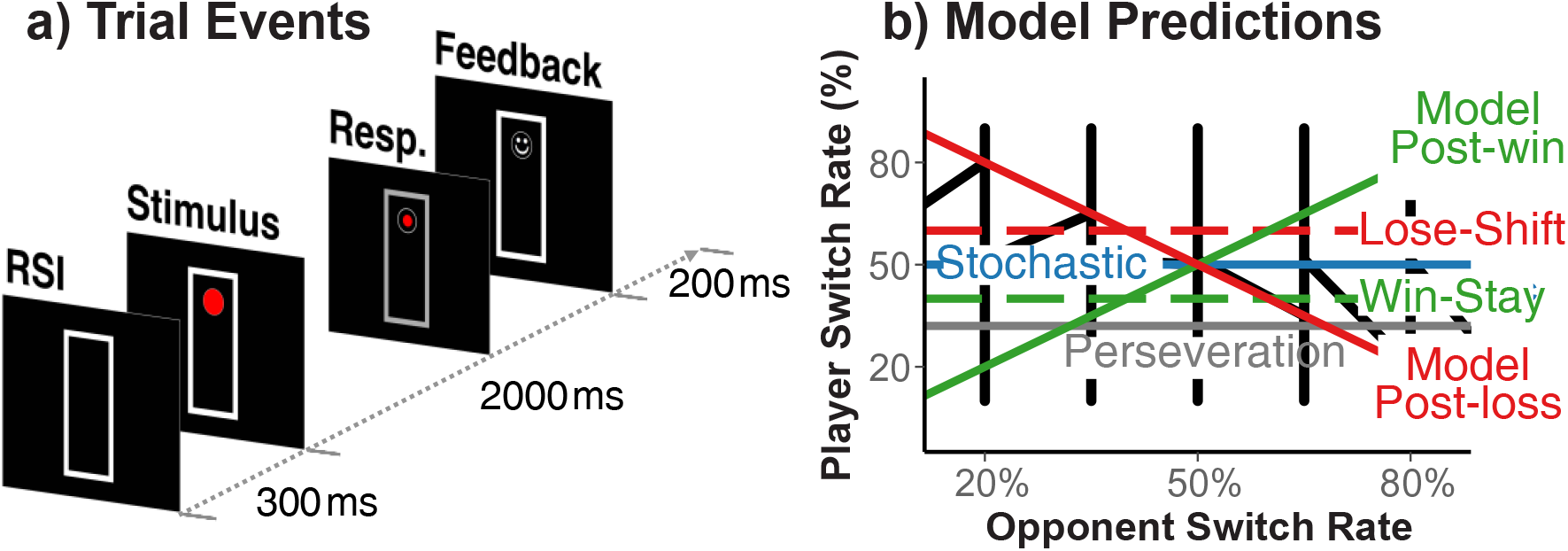
a) Sequence of trial events and response rules in the fox/rabbit paradigm. b) Idealized predictions of how different choice strategies and biases are expressed in the player’s switch rate as a function of the opponent’s switch rate. Choices based on an internal model of the opponent, lead to a positive relationship between the player’s switch rate following wins (green filled line) and to a negative relationship following losses (red filled line). Memory-free, stochastic choice leads to a 50% switch rate irrespective of the opponent’s switch rate (blue line). The hypothesis of model-based choice after wins and stochastic choice after losses predicts the combination between the filled green and the blue line. A perseveration bias leads to an unconditional reduction of the switch rate (gray line) and a win-stay/lose-shift tendency to a selective increase following loss trials and a decrease following win trials (dashed green and red lines).

In order to establish the degree to which players utilized a model of their opponent, we exposed them to a set of simulated opponents that differed in their average switch rate (e.g., 20%, 35%, 50%, 65%, and 80%), but otherwise behaved randomly. Variations in opponents’ switch rate provide a diagnostic indicator of both model-based and stochastic behavior (Figure 1b). Specifically, a model-based agent should appreciate the fact that when playing against an opponent who switches frequently between run and freeze moves (i.e., *p*>.5), it is best to switch moves after a win (i.e., “following along with the opponent”), but to stick with the same move after a loss (i.e., “waiting for the opponent to come to you”); the opposite holds for opponents with a low switch rate (i.e., *p*<.5). Thus, model-based behavior would produce a combination of the filled green and red switch-rate functions in Figure 1b. In contrast, a memory-free agent should produce random behavior (i.e., a switch rate close to *p*=.5) irrespective of the opponent’s strategies (i.e., the blue function in Figure 1b). Thus, our hypothesis of a feedback-contingent mix between model-based and stochastic behavior predicts an increase of players’ switch rate as a function of their opponents’ switch rate for post-win trials (the filled green function in Figure 1b), but a switch rate close to *p*=.5 irrespective of the opponent’s switch rate on post-loss trials (the blue function in Figure 1b).

In standard, sequential choice paradigms, people choose between simple actions. In contrast, the fox/rabbit task requires choices between two different action rules. The rules, combined with the stimulus determine simple actions that need to be executed accurately and under time pressure. This procedure has the advantage of allowing a distinction between two potential sources of stochasticity. Stochasticity may simply be the result of an increase of unspecific information-processing noise in the system, which would not only lead to more random choices between action rules, also more error-prone and slower action selection. Alternatively, choice stochasticity may be specifically due to reduced choice input from model/context representations. In this case, an increase in choice stochasticity should not be associated with more error-prone and slower action selection.

### Analytic Strategy for Testing Main Prediction

To test the prediction of loss-induced stochastic behavior, we cannot simply compare the slopes of post-win and post-loss switch-rate functions. Such a comparison would not differentiate between a pattern of post-loss and post-win functions with the same slope but opposite signs (as would be consistent with the model-based choice strategy, see Figure 1b) and the predicted pattern of more shallow slopes following losses. Therefore, as a general strategy, we tested our main prediction by comparing slopes after selectively inverting the labels for the opponent switch-rate in the post-loss condition (e.g., 80% becomes 20%). This allows direct comparisons of the steepness of post-win and post-loss switch-rate functions. In the SI, we also present results from standard analyses.

### Modeling Choice Behavior

Our behavioral indicator of a mix between model-based and stochastic behavior is expressed in players’ switch rate, which can also be affected by the perseveration and win-stay/lose-shift bias (see Figure 1B). In standard sequential-decision paradigms it is very difficult to distinguish between stochastic behavior and low-level biases. Therefore, we attempted to obtain a realistic characterization of the various influences on choice behavior by using simple choice model to predict the probability of switch choices *p*_switch_:

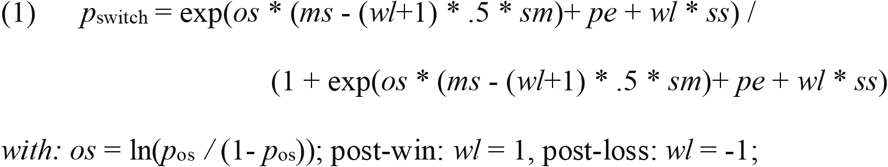

where *p_os_* is the opponent’s switch rate, which is translated into its log-odds form (*os*); *wl* codes for wins versus losses on trial *n*-1. The parameter *ms* (**m**odel **s**trength) represents the strength of the internal model (*m*s=1 would indicate direct probability matching between the opponent’s and the player’s switch probability). The parameter *sm* (**s**trategy **m**ix) represents the degree to which the model-based choice is changed on post-loss relative to post-win trials; a negative *sm* parameter would indicate suppression of the model in favor of stochastic choice following losses. In addition, a positive *pe* (**pe**rsevertion **e**ffect) parameter represents the tendency to unconditionally favor the previously chosen task, and a positive *ss* (win-**s**tay/lose-**s**hift) parameter expresses the strength of the win-stay/lose-shift bias. We present predictions from this model in Figure 2, and report additional details of the modeling results in the SI.

**Fig. 2.**
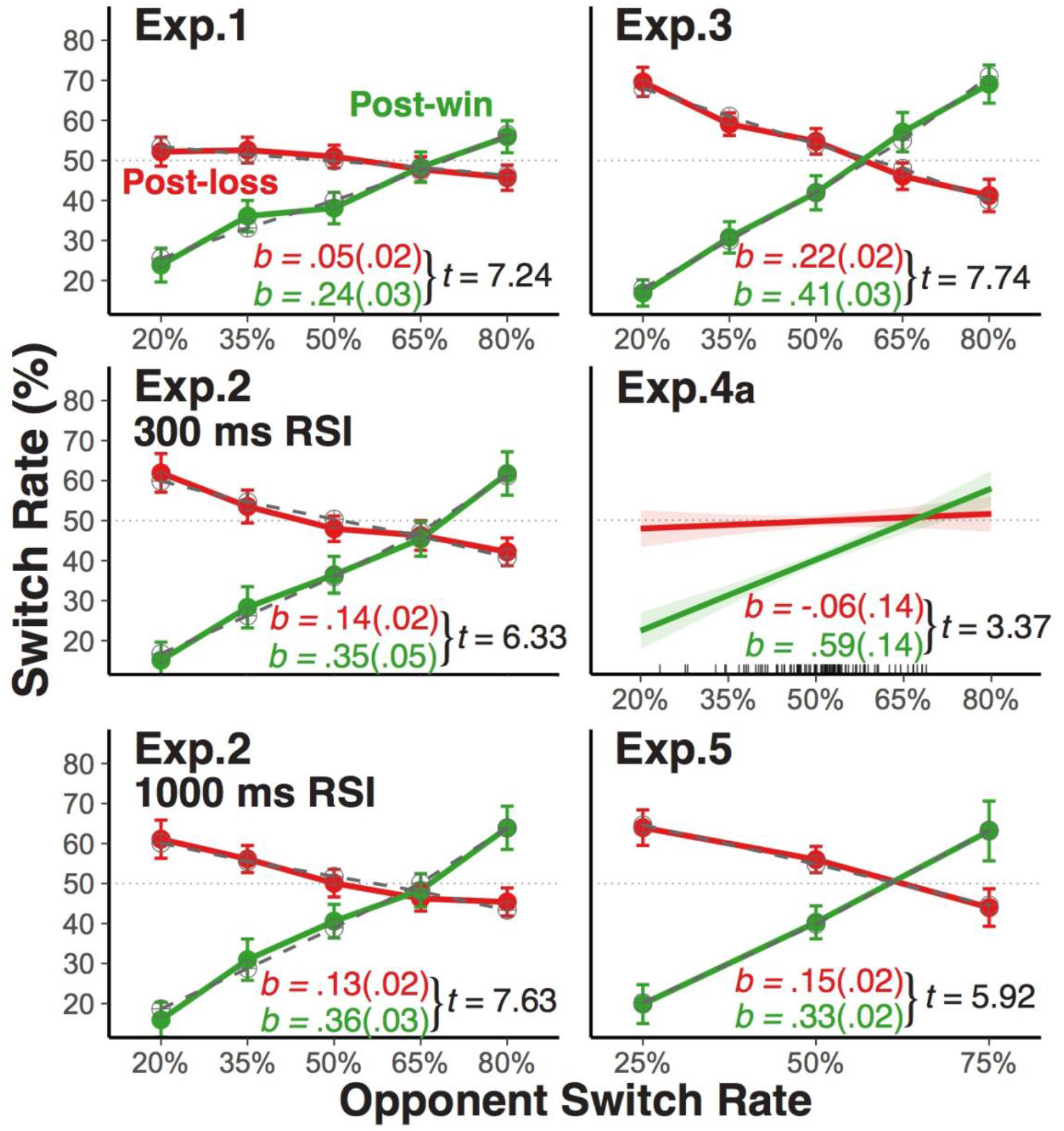
Average empirical switch rates for post-win and post-loss trials as a function of the simulated opponents’ switch rates for Experiments 1, 2, 3, and 5 and the average switch rate of each human opponent in Experiment 4a (tick marks on x-axis indicate individual averages). The dashed lines for Experiments 1,2, 3, and 5 show the predictions of the theoretical choice model applied to the group average data (see text and Supplemental Information for details). Error bars represent 95% within-subject confidence intervals. For the analyses, we regressed the player’s switch rate on the opponents’ switch rates, the win-loss contrast, and the interaction between these two predictors after reversing the labels of the opponents’ switch-rate predictor for post-loss trials (see section *Analytic Strategy for Testing Main Prediction*). As a test of these interactions, we show the corresponding *t*-values (*SE*); the unstandardized slope coefficients (*SE*; green=post-win, red=post-loss) were derived from separate analyses for post-win and post-loss trials.

### Choice Behavior with Simulated Opponents

Experiment 1 establishes the basic paradigm. Figure 2 (Exp. 1) shows that participants increased their switch rate as a function of their opponents’ switch rates following win trials. In contrast, on post-loss trials, the slope of the function relating participants’ switch rate to their opponents’ switch rate was although slightly negative, much smaller than on post-win trials and it was centered at *p*=.5, a pattern that is consistent with largely stochastic choice. The condition with an opponent switch-rate of *p*=.5 most closely resembles previous studies that have reported a win-stay/lose-shift bias in competitive situations (10). In fact, for this condition, we did find a significantly higher switch rate after losses than after wins, indicating that reinforcement-based tendencies are one factor that affects choice. Likely, win-stay/lose-shift and perseveratory tendencies are also responsible for participants’ apparent reluctance to fully endorse the model-based strategy after wins, as indicated by the fact that the slope of the switch-rate function after wins is substantially smaller than 1. Indeed, here and in the remaining experiments, results from applying our choice model to the data indicate that (a) a strong tendency towards model-based choices on post-win trials, (b) an increase of stochastic choice on post-loss trials, (c) a general perseveratory tendency, and (d) a win-stay/lose-shift bias all contribute to the overall choice behavior (see Supplemental Information).

After feedback from the previous trial, participants had only 300 ms to choose their move for the next trial in Experiment 1. Therefore, one might argue that the observed stochastic choice is simply a result of negative feedback temporarily interfering with model-based selection (16). To examine this possibility, we manipulated the response-to-stimulus interval (RSI) in Experiment 2 between 300 ms and 1000 ms. As shown in Figure 2, this manipulation had no effect, indicating that stochastic choice is not due to loss-induced processing constraints.

The fox/rabbit task was modeled after the voluntary task-switching paradigm in order to recreate executive control demands of actual competitive situations and to allow a separation between choice stochasticity and more general increase of noise in the cognitive system. However, it is important to explore how the observed pattern might change with less complex response rules than used in this paradigm. We therefore implemented in Experiment 3 simple choices without any contingencies on external inputs (i.e., the fox wins when selecting the same up or down location as the rabbit, and vice versa). Here, we generally found a stronger expression of model-based choice following *both* losses and wins, and also much less perseveration bias (see Supplemental Information for details). Yet the win-loss difference in slopes remained just as robust as in the other experiments (Figure 2). Thus, the more complex actions that players had to choose from in Experiments 1 and 2 may have suppressed the overall degree of model-based action selection (Otto et al, 2014). However, response rule complexity did not appear to affect the win/loss-contingent difference in the relative emphasis on model-based versus stochastic choices.

### Competition against Human Players

It is possible that the observed pattern of results is specific to experimental situations with a strong variation in simulated, opponent switch rates. To examine the degree to which this pattern generalizes to a more natural, competitive situation, we used in Experiment 4a pairs of participants who competed with each other in real time, with one player of each dyad acting as fox, the other as rabbit (see also Supplemental Information, for Experiment 4b as a non-competition control experiment). Obviously, the naturally occurring variation in switch rates was much lower here (see distributions of individuals average switch rates in Figure 2). Nevertheless, the estimated slopes linking players’ switch rates to opponents’ switch rates exhibited a very similar pattern as for the simulated-opponent experiments. In addition, a trial-by-trial version of our choice model revealed an overall pattern that was qualitatively consistent with the results from the simulated-opponent experiments (see Supplemental Information for details).

### RT and Error Effects

Different from standard choice paradigms (23, 24), the current paradigm allows us to distinguish between stochasticity during the choice between action rules and general information-processing noise (25). If the loss-induced choice stochasticity is due to a general increase in information processing noise then we should see that greater stochasticity goes along with more erroneous and/or slower actions. Figure 3 shows each individual’s degree of model-based choice (expressed in terms of absolute switch-rate slopes) after loss and win trials and as a function of both RTs or error rates. In most experiments, there is a slight increase in error rates following loss trials. However, across individuals, the substantial reduction in model-based choice after loss trials is *not* associated with a consistent increase in error rates or RTs.

**Fig. 3.**
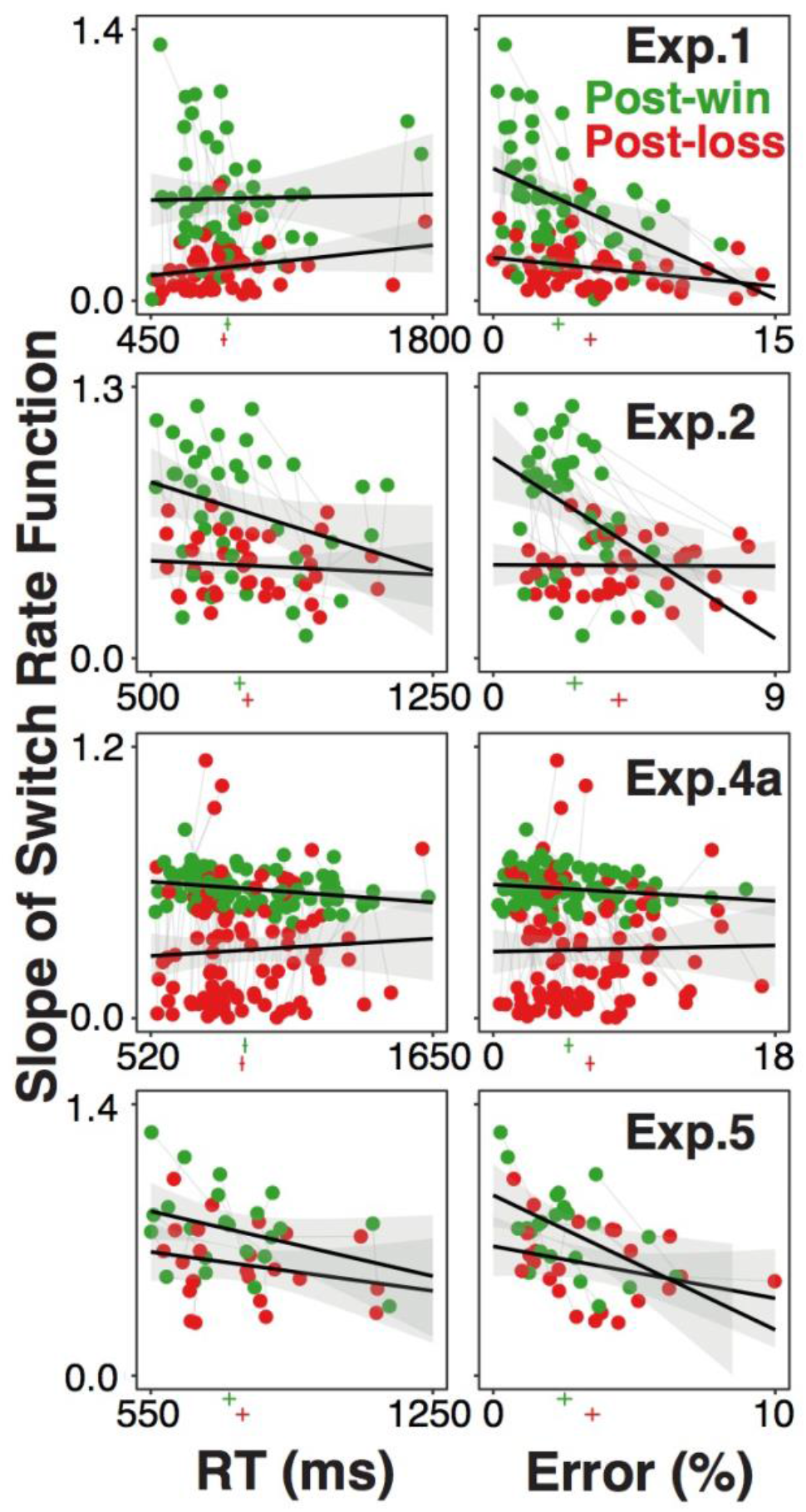
Individual participants’ degree of model-based choice (indicated by the slopes of switch-rate functions) in relationship to RTs and error rates, separately for post-win (green) and post-loss trials (red), and for each experiment using the rule-selection paradigm. Each participant is represented both in the post-win and the post-loss condition. The green and red vertical lines below the x-axis of each graph indicate average RTs and error rates, the horizontal line the corresponding 95% confidence interval (within subject). If the increase of choice stochasticity between post-win and post-loss trials were due to greater, general information-processing noise, then the win/loss-related decrease in slopes of the switch-rate functions would be accompanied by consistent increases in RTs and/or error rates.

In Figure 3, this is evident in the large portion of individuals with differences in model-based behavior as a function of post-win and loss trials, but with very similar error rates or RTs. Likewise, in multilevel regression models with the absolute switch-rate slopes as dependent variable, the post-win/loss contrast remained highly robust after controlling for RTs and errors as within-subject fixed effects (range of *t*-values associated with the post-win/loss predictor: 3.96-10.78).

### Loss-induced Increase of Stochastic Choice

So far, we have established that participants were more sensitive to their opponents’ global strategies (i.e., the average switch rates) following win than following loss trials. Next, we examined the degree to which these win-loss differences generalized to players’ consideration of the recent history of their opponents’ and their own choices. To this end, we used multi-level logistic regression models with the switch/repeat choice as criterion. The models included the trial *n*-1 to *n*-3 switch/repeat decisions for opponents and for players, along with the opponents’ overall switch rate and were separately run for post-loss and post-win trials to generate the coefficients presented in Figure 4. To directly compare the size of the coefficients, irrespective of their sign, we again reversed the labels, both for the opponents’ global switch rate, but also for both the opponent’s and the player’s *n*-1 to *n*-3 switch/repeat decisions (e.g., switch becomes repeat; see section *Analytic Strategy for Testing Main Prediction* and *History Analyses* in the Supplemental Information). Consistent with the prediction that switch/repeat choices following losses are less dependent on recent history, the coefficients for the opponents’ history and also the players’ own history were in most cases substantially lower after loss than after win trials.

**Figure 4.**
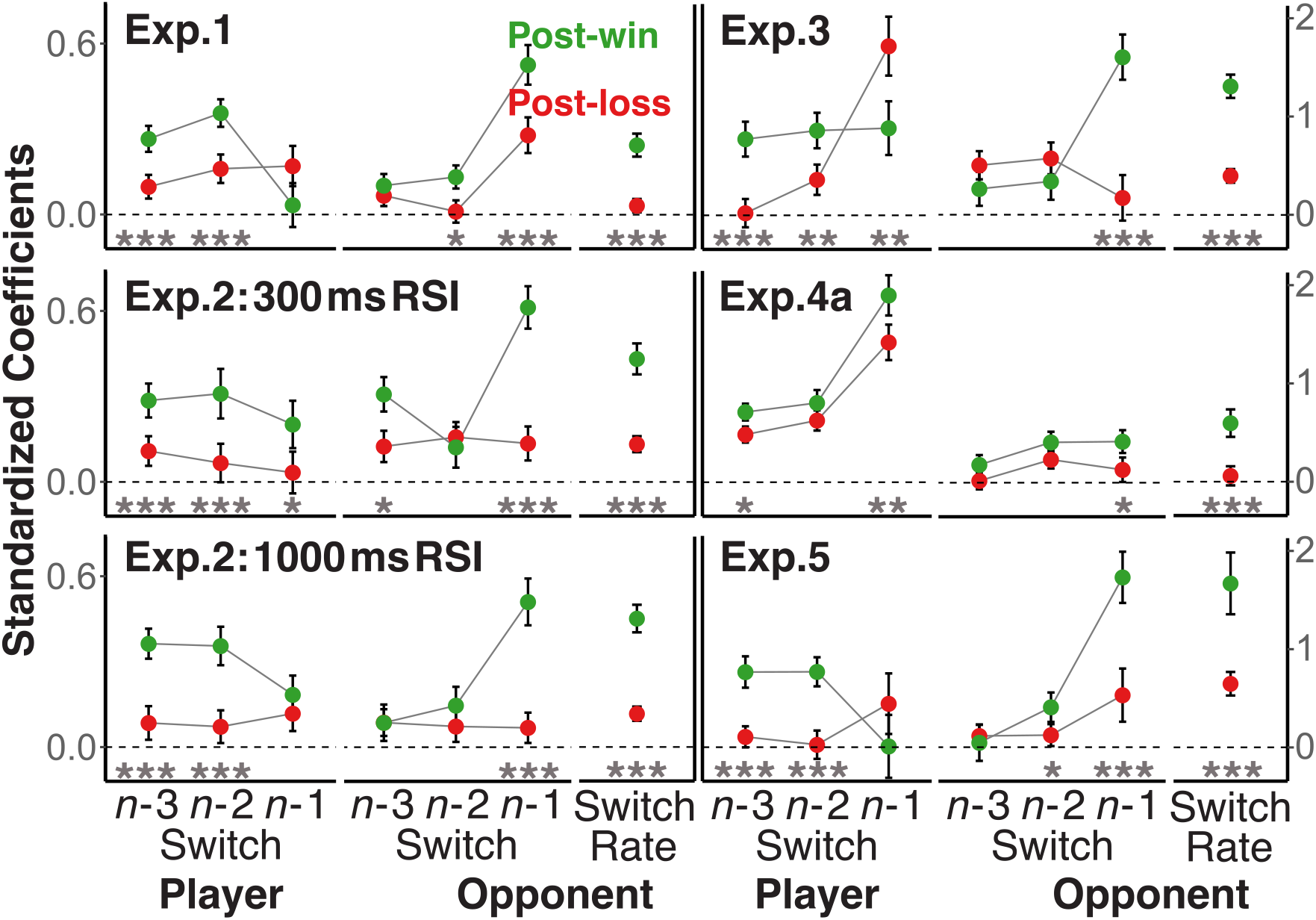
Standardized coefficients from multi-level logistic regression models predicting the trial *n* switch/no-switch choice on the basis of players’ and opponents’ switch/repeat choices on trials *n*-1 to *n*-3 and the opponents’ overall switch rate. Error bars are within-subject, standard errors around the coefficients. To focus on the difference in the strength of relationships rather than their sign, the labels for all opponent-related predictors were reversed for post-loss trials (see section *Analytic Strategy for Testing Main Prediction*). In addition, we also reversed the labels for all player-related predictors with a win/loss switch in sign (for signed coefficients, see Figure S6). For a statistical test of the size difference between post-win and post-loss coefficients, all these history/context variables were included into one model together with the post-win/post-loss contrast and the interaction between this contrast and each of the history/context predictors. Significance levels of the interaction terms are indicated in the figure, *<.05, **<.01, ***<.001.

### Neural Evidence for Memory-Free Choice Following Losses

Research with animal models and human, neuroimaging work indicates that midfrontal brain regions, such as the anterior cingulate cortex are involved in action-relevant representations and in the gating between different modes of action selection (26–28). Further, a large body of research suggests that midfrontal EEG activity in response to action feedback contains prediction error signals (10, 29–33), which in turn are reflective of action-relevant expectancies (i.e., the current task model). Therefore, it is theoretically important to link our behavioral results to this broader literature. Specifically, it would be useful to show that (a) only on post-win, but not on post-loss trials, the midfrontal EEG signal contains information about the choice context/model, and (b) that the context information contained in the EEG signal is in fact predictive of upcoming choices.

In Experiment 5, we assessed EEG while participants played the fox/rabbit game against three different types of opponents (25%, 50%, 75% switch rate; see Figures 2 and 3 for behavioral results). We conducted a two-step analysis. In the first step, we tested the prediction that the mid-frontal EEG signal contains less information about the choice-relevant context after loss-feedback than after win-feedback. To this end, we regressed trial-to-trial EEG signals on A) the opponent’s overall switch rate, B) the opponent’s lag-1 switch/no-switch, C) the player’s lag-1 switch/no-switch, and D) the interaction between A) and B), that is between the local and global switch expectancies. The latter term was included to capture the fact that if feedback-related EEG reflects expectancies about opponents’ switch rates, local switch expectancies may depend on the global switch-rate context (31).

The standardized coefficients shown in Figure 5a (see caption for details) indicate the amount of information about each of the four context variables that is contained in the mid-frontal EEG signal. As apparent, the EEG signal showed a robust expression of the history/context variables following win feedback. Following loss feedback, context information is initially activated, but then appears to be suppressed compared to post-win trials, and trends towards zero at the end of the feedback period. Accordingly, coefficients were significantly larger in post-win trials than in post-loss trials, opponents’ overall switch rate: *b*=0.07, *se*=0.01, *t*(25)=5.22, *p*<.001, opponents’ lag-1 switch/no-switch: *b*=0.04, *se*=0.01, *t*(25)=4.14, *p*=.001, player’s lag-1 switch/no-switch: *b* =0.01, *se*=0.009, *t*(25)=1.19, interaction between opponents’ overall switch rate and lag-1 switch: *b* =-0.08, *se*=0.01, *t*(25)=-7.22, *p*<.001. Given that feedback is related to subject’s propensity of switching on the upcoming trial, it is in principle possible that these coefficients simply reflect preparation or increased effort for the upcoming switch. However, as we show in the Figure S7, controlling for upcoming switches has negligible effects on the results presented in Figure 5a.

**Fig. 5.**
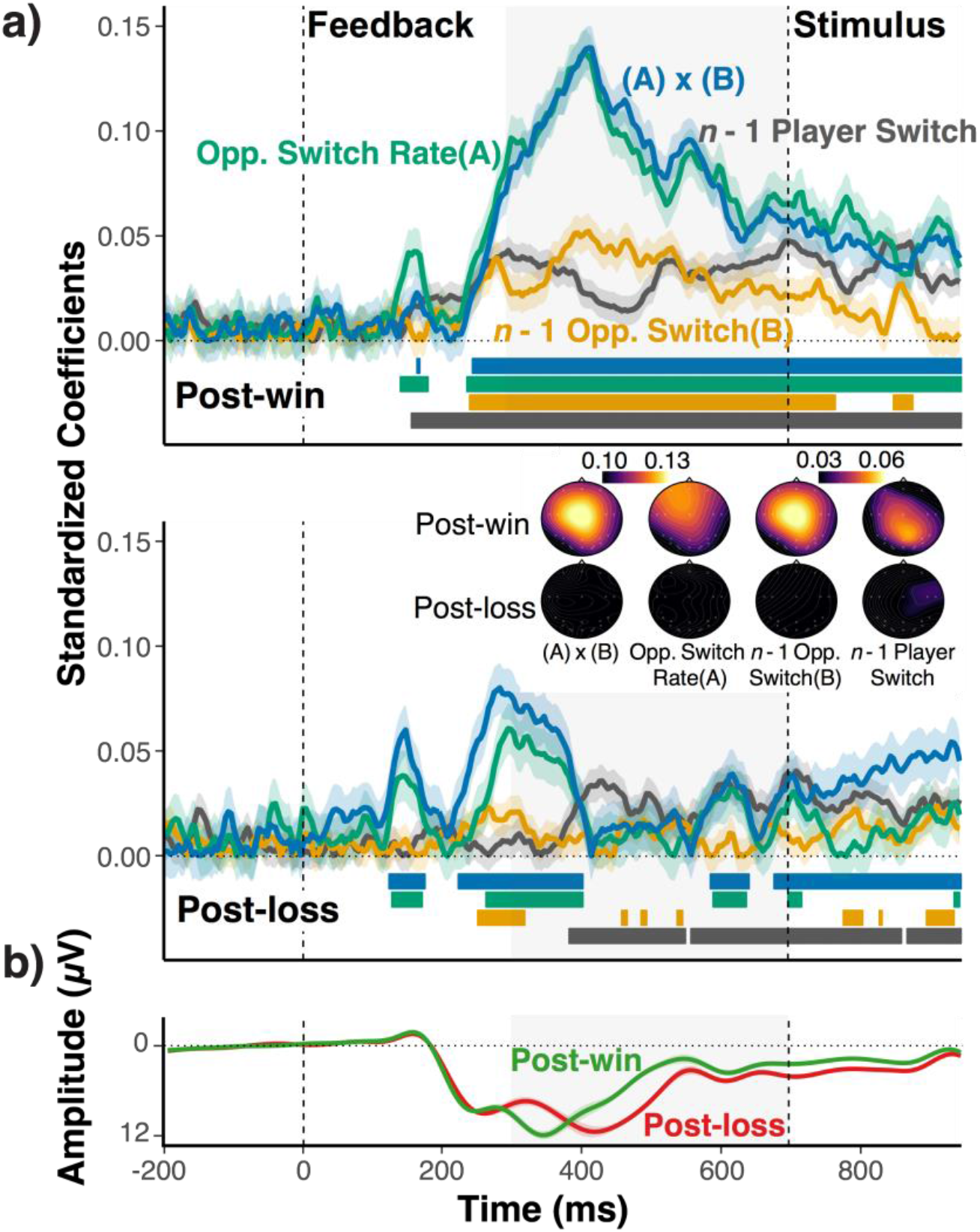
a) Standardized coefficients from multi-level regression models relating EEG activity at Fz and Cz electrodes to the opponent’s overall switch rate (A), the *n*-1 opponent switch/no-switch choice (B), the *n*-1 player’s switch/no-switch choice, and the interaction between A) and B) for each time point and separately for post-win (upper panel) and post-loss (lower panel) trials. Shaded areas around each line indicate within-subject standard errors around coefficients. As coefficients for opponent-related predictors showed a marked, win/loss flip in sign, we again reversed the labels of the post-loss predictors (see section *Strategy for Testing Main Prediction* and Figures 2 and 3; for signed coefficients, see Figure S10). For illustrative purposes, colored bars at the bottom of each panel indicate the time points for which the coefficients were significantly different from zero (*p* < .05). See text for statistical tests of the predicted differences between coefficients for post-win and post-loss trials. The insert shows the topographic maps of coefficients that result from fitting the same model for each electrode separately. b) Average ERPs for post-win and post-loss trials, showing the standard, feedback-related wave form, including the feedback-related negativity (i.e., the early, negative deflection on post-loss trials). Detailed ERP results are presented in the Supplemental Information (see Figure S8).

Our analytic strategy deviates from the standard approach of analyzing the EEG signal in terms of feedback-locked, event-related potentials (ERPs; see Figure 5b). We used our approach because we did not have a-priori predictions about how exactly the combination of different history/context variables would affect ERPs. More importantly, our regression-based approach naturally yields trial-by-trial indicators of the expression of context-specific information, which can be used in the second step of our analysis (see below), and which would be difficult to obtain through standard ERP analyses. In the Supplemental Information (Figure S6) we also show that the ERP results are indeed generally consistent with a prediction-error signal that is more strongly modulated by the choice context after wins than losses.

The conclusion that post-loss stochastic behavior occurs because context representations are suppressed, would be further strengthened by evidence that the information contained in the EEG signal is actually relevant for upcoming choices. Therefore, as the second step, we conducted a psychophysical interaction (PPI) analysis (34). As only the post-win trials showed robust context information in the EEG signal, we focus here on these trials (but see Table S4 for an analysis of all trials). In a multi-level, logistic regression analysis, we predicted players’ trial *n* switch choices, based on (1) the trial *n*-1 residuals from the preceding analysis (reflecting trial-by-trial variations in the EEG signal after controlling for the four context variables), (2) the set of four context variables from the preceding analysis for trial *n*-1, (3) and the corresponding four interactions between the residuals and the context variables. We found significant main effects for the residual EEG signal and all context variables (see Table S4). Most importantly, the residual EEG signal modulated how the upcoming choice was affected by the opponent’s lag-1 switch/repeat: *b*=-0.67, *se*=0.05, *t*(25)=-3.31, *p*<.01, and the opponent’s overall switch rate: *b* = 0.13, *se*=0.06, *t*(25)=2.26, *p*<.05. These results indicate that the information about context variables contained in the EEG signal is indeed relevant for choices.

As a final step, we also examined to what degree variations in the strength of history/context representations can account for individual differences in choice behavior. To this end, we derived for each individual and predictor, the average, standardized coefficient from the analysis presented in Figure 5 across the 300 ms to 700 ms interval. Separately for post-win and post-loss trials, we correlated these scores with two behavioral measurements: 1) individuals’ switch-rate functions as an indicator for model-based choice and 2) the overall rate of winning. For post-loss trials, we again used opponent-related predictors with reversed labels (see EEG Analysis section for details). Thus, for all analyses, more positive scores are indicative of individuals with more model-conform behavior.

As shown in Figure 6, coefficients from post-win EEG signals generally predicted the variability among individuals in the degree of model-based adaptation and the rate of winning (except for coefficients of *n*-1 player’s switch). In contrast, such relationships were absent for post-loss trials. We also tested the difference between post-win and post-loss correlations using a *z*-test for dependent correlations with either non-overlapping (for switch-rate function slopes) or overlapping samples (for overall rate of winning). For the switch-rate function slopes, we found significant differences for opponents’ lag-1 switch/no-switch, *z*(25)=2.87, *p*=.003, and the interaction between opponent’s overall switch rate and lag-1 switch/no-switch choice, *z*(25)=4.26, *p*<.001, but not for opponent’s overall switch rate, *z*(25)=1.38, *p*=.16 and the player’s lag-1 switch/no-switch, *z*(25)=- 0.76, *p*=.45. Similarly, for the overall rate of winning, we found significant differences for opponents’ lag-1 switch/no-switch choice, *z*(25)=2.33, *p*=.02, and the interaction between opponents’ overall switch rate and the lag-1 switch/no-switch, *z*(25)=2.21, *p*=.02, but again not for opponents’ overall switch rate, *z*(25)=1.54, *p*=.12, and the player’s lag-1 switch/no-switch, *z*(25)=-1.18, *p*=.23.

**Fig. 6.**
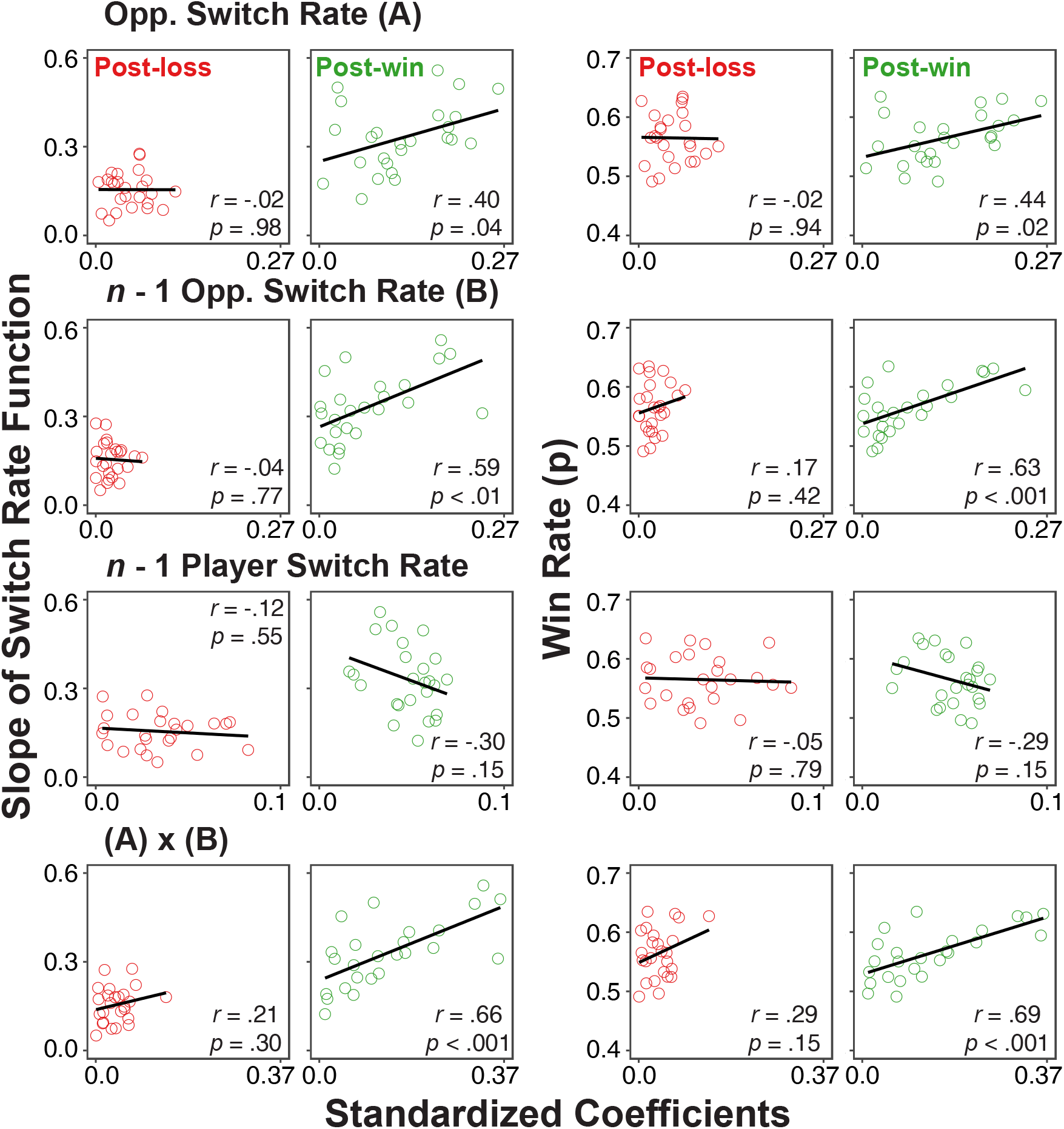
Correlations between individuals’ standardized coefficients from the multi-level regression analysis relating the EEG signal to the different history/context variables and 1) their slopes for the switch-rate functions (left two columns) or 2) their overall win rate (right two columns) separately for post-win and post-loss conditions. Coefficients were obtained by fitting models with the EEG signals averaged over a 300-700 ms interval of the post-feedback period (the shaded interval in Figure 5).

Combined, these individual differences results suggest that the degree to which history/context variables are represented in the EEG signal following win feedback, predicts both individuals’ reliance on the model of the opponent and their overall competitive success. Consistent with the idea that following loss-feedback, model-based representations are suppressed, these relationships are largely absent on post-loss trials. With its relatively small sample size, this experiment was not designed as an individual differences study and therefore these exploratory results need to be considered with caution. However, confidence in these results is strengthened by the fact that they are consistent with the findings from the within-subject PPI analyses.

## Discussion

Our results show that people can use two different choice regimes for selecting their next move in a competitive game. Immediately following a win, participants tended to rely on an internal model of the opponent’s behavior and his/her general tendency to switch moves relative to the preceding trial. Following a loss trial, they selected their next move more stochastically and less influenced by the local or global choice context (15, 22). At least in humans, the demonstration that after loss feedback, model-based selection is largely replaced by a more memory-free, stochastic mode of selection is a theoretically important result.

Past work has characterized behavior in zero-sum game situations as a problem of a trade-off between model-free, win-stay/lose-shift tendencies and choices based on a model of the opponent (e.g., 14). In addition, work with the voluntary task-switching paradigm has suggested that people generally have a tendency to repeat action plans that were executed in the immediate past (8, 9). Indeed, our modeling results show that each of these influences is consistently present in our data. Importantly, the tendency towards model-based choice and the tendency towards stochasticity on post-loss trials, independently predicted individuals’ competitive success, and over and above the effect of the known, lower-level biases (see Table S2 and S3).

Our findings are generally consistent with recent research indicating an increase of decision noise––defined as deviations from optimal choice––when exploration is beneficial in a sequential decision situation (35). However, it is a novel finding that a stochastic model of selection is turned off and on according to positive versus negative feedback. Also, different than in standard sequential-decision paradigms, in our paradigm subjects chose action rules rather than specific actions (except for Experiment 3), allowing us to separately examine choices and the efficiency of action execution. The fact that the post-loss increase in stochastic choice was *not* accompanied by a consistent increase in action errors or RTs, speaks against the possibility that choice stochasticity results from a general increase in information-processing noise.

The results we report here are reminiscent of an emerging literature on the asymmetric consequences of positive versus negative feedback(36, 37). Specifically, people appear to update their beliefs to a lesser degree following negative feedback than positive feedback, and this asymmetry may be responsible for a generalized optimism bias. However, it is also important not to confuse our results with these existing findings. The asymmetric updating rate affects representations relevant for the action choice that *preceded* the positive or negative feedback. In contrast, we report here that following negative feedback, people make less use of relevant representations as they choose the *next* action. Combined, these sets of findings suggest that negative feedback negatively affects both learning, and the use of existing information. The degree to which these two biases share a common, underlying mechanism is an important question for further research.

In many competitive situations, a model of the opponent is needed to exploit regularities in the opponent’s behavior (1, 4). At the same time, one’s own choices need to appear unpredictable to the opponent. The feedback-contingent mix of choice regimes we report here, may be an attempt to meet the opposing demands within the limitations of our cognitive system. By this account, wins signal to the system that the current model is valid and is safe to use. In contrast, losses signal that the current model may be invalid and that alternatives should be explored and/or that there is a danger of being exploited by the opponent. As a result, the current model is temporarily abandoned in favor of stochastic, memory-free choice.

Viewed by itself, the turn towards stochastic choice following losses is an irrational bias. Indeed, our modeling results indicate that the degree to which players switch to stochastic choice after losses, negatively predicts their success in competing against both simulated and actual opponents (see Tables S2 and S3). Interestingly, this choice regime also resembles maladaptive, learned-helplessness patterns that are typically observed across longer time scales and that are often associated with the development of depressive symptoms (38, 39). To what degree the trial-by-trial phenomenon examined here and the longer-term, more standard learned-helplessness processes are related is an interesting question for further research. It is however also important to consider the possibility that the loss-contingent switch to stochastic choice is adaptive in certain circumstances. For example, in situations with a greater number of choices or strategies than were available within the fox/rabbit game, a switch to stochastic choice may allow the exploration of neglected regions of the task space (29). Also, given known constraints on consistent use of model-based selection, the ability to revert to stochastic behavior provides a “safe default” that at the very least reduces the danger of counter-prediction through a strong opponent.

In this regard, a recent study by Tervo at al. (15) is highly relevant. These authors analyzed the choice behavior of rats playing a matching-pennies games against simulated competitors of varying strength. The animals showed model-based choice behavior against moderately strong competitors, but switched to a stochastic choice regime when facing a strong competitor. The switch in choice regimes occurred on different time scales in the Tervo et al. study (i.e., several sessions for each competitor) than in the current work (i.e., trial-to-trial). Nevertheless, it is remarkable that a qualitatively similar, failure-contingent switch between choice regimes could be found in both rats and human players.

Tervo et al. also used circuit interruptions in transgenic rats to show that a switch to stochastic choice is controlled via noradrenergic input to the anterior cingulate cortex (ACC), which supposedly suppresses or perturbs ACC-based representations of the current task model. Interestingly, in our Experiment 5, we found that EEG signals registered at mid-frontal electrodes, contained robust information about the opponents’ and the players’ own strategies following win-feedback. On post-loss trials the EEG signal initially contained information about the opponent’s global and local behavior, but this information was all but eliminated by about 400 ms following the feedback signal. This time-course suggests that context information is available in principle, but is quickly suppressed on post-loss trials. Additional analyses indicated that the task-relevant information contained in the EEG signal was indeed relevant for upcoming choice behavior. Feedback-contingent, mid-frontal EEG signals are often thought to originate in the ACC and associated areas (22, 31, 40). Thus, our results are fully consistent with the theory that these brain areas are critical for representing the current choice-relevant information. Further, the noradrenergic perturbation process identified by Tervo et al., suggests an interesting hypothesis for future research about how in humans, task-relevant representations might be actively suppressed to promote memory-free, stochastic choice (41). More generally, this emerging body of evidence provides one possible answer to the fundamental question how a memory-based choice system can produce non-deterministic behavior––namely through selectively “losing” or suppressing memory records of the choice context.

## Methods

### Participants

Subjects were University of Oregon students who participated after giving informed consent in exchange for monetary payment or course credits; Experiment 1: *N*=56 (38 female), Experiment 2: *N*=40 (28 female), Experiment 3: *N*=44 (25 female), Experiment 4a: *N*=100 (62 female), Experiment 4b: *N*=40 (22 female; presented only in SI), Experiment 5: *N*=25 (13 female). Four subjects from Experiment 1 and three pairs from Experiment 4a were excluded, because the experimental session could not be completed. The entire study protocol was approved by the University of Oregon’s Human Subjects Review Board (Protocol 10272010.016).

### Stimuli, Tasks and Procedure

On each trial of the fox/rabbit game, players observed a circle either on the bottom or the top of a vertically aligned rectangle. They had to choose between one of two rules for responding to the circle location. The “freeze rule” implied that the circle stayed at the same location and it required participants to press among two keys the one that was compatible with the circle location (‘1’ and ‘4’ on the number pad). The “run rule” implied that the circle moved to the opposite location within the vertical box and participants had to press among a separate set of vertically aligned keys (‘5’ and ‘8’ on the number pad) the key that was incompatible with the circle location (9). On a given trial, the fox player won 2 cent per trial, when both players chose the same rule, whereas the rabbit player won when choices were different. Participants had to respond within a 2000 ms interval and after that interval, they received feedback presented for 200 ms with a smiley face indicating a win trial and a frowny face a loss trial. Any erroneous responses were followed by a frowny face. For Experiment 1, 3, and 4, the inter-trial interval (ITI) was 300 ms. For Experiment 2, the ITI was randomly selected to be 300 ms or 1000 ms on trial-by-trial basis and for Experiment 5 the ITI was 750 ms to allow assessment of feedback-related EEG activity. For Experiment 3 (“simple response”), each trial was initiated by a circle appearing at the center of the vertically arranged stimulus rectangle. Using the vertically arranged “1” or “4” keys, participants had to shift the circle up or down within the rectangle. Again, matching moves between opponents implied a win for the fox and a loss for the rabbit player.

For all experiments except for Experiments 4a and 4b, participants faced a variety of simulated opponents that differed in terms of switch-rate strategies. Participants were instructed that the different simulated players represented common strategies that one might find in human players. At the beginning of each block, participants were notified that they would be facing a new, simulated opponent, and whether they played the role of the fox or the rabbit, but received no instruction about the specific strategies. For Experiment 1, 2, and 3, switch rates varied on a block-by-block basis between 20%, 35%, 50%, 65%, and 80% randomly. For Experiment 5, 25%, 50% and 75% switch rates were used. In Experiments 1, 2, and 3, participants worked on 10 blocks with 80 trials each (i.e., 5 opponent strategies x fox/rabbit roles) and with 24 blocks of 80 trials in Experiment 5 (i.e., 4 repetitions of 3 opponent strategies x fox/rabbit roles). In Experiment 4a participants were paired into fox/rabbit dyads and played in real-time on two computers within the same room, but without opportunity for direct communication. Here participants played 7 blocks of 80 trials each. Experiment 4b served as a non-competition control experiment that was otherwise identical to Experiment 4a. The only difference was that here trial-by-trial wins and losses were completely random and participants were informed that this was the case. All experiments started with 80 practice trials without a competitor in order to familiarize participants with response procedures. The experiments were programmed in Matlab (Mathworks) using the Psychophysics Toolbox(42) and presented on a 17-inch CRT monitor (refresh rate: 60Hz) at a viewing distance of 100 cm.

### EEG Recordings

In Experiment 5, Electroencephalographic (EEG) activity was recorded via 20 electrodes and processed using standard procedures (see Supplemental Information for details). Single trial EEG signals were segmented into 1250 ms epochs starting from 200 ms before the onset of feedback. Thus, each epoch included 700 ms post-feedback periods and the initial 250 ms intervals of the next trials. Each electrode’s EEG signal was also pre-whitened by linear and quadratic trends across experimental trials and blocks. After baselining signals with data from the initial, 200 ms interval, EEG activity from electrodes Fz and Cz, was averaged. These electrodes were selected based on previous studies reporting a robust interaction between the feedback and the probability context during reinforcement learning (43). The resulting signal was regressed via multilevel modeling with two levels (i.e., trials nested within participants) on context variables, as described in the Results section. For illustrative purposes, this was done on a time-point by time-point basis (see Figure 4). To conduct statistical tests of the post-win versus post-loss regression coefficients for the psychophysiological interaction analysis predicting choices, for the individual differences (Figure S11), and for the topographic maps (Figure 4), we averaged the EEG signal for an a-priori defined 300-700 ms interval following the feedback signal. This interval is based on the typical time-course of feedback effects reported in the literature (43). The difference between post-win/loss models was tested in the same manner as in the multilevel model for history effects, namely by inverting predictors of opponents’ history/context for post-loss trials (see section *Analytic Strategy for Testing Main Prediction* and *History Effects Analysis* in the Supplemental Information)

### Data Availability Statement

The data that support this study, along with the analysis scripts are available through the Open Science Foundation repository (https://osf.io/j6beq/).

## Acknowledgements

This work was supported in part by National Instute of Aging grant R01 AG037564-01A1

## SUPPLEMENTAL INFORMATION

### Additional Information on Analyses and Methods

#### History Analyses

To evaluate the predictability of the current choice by the recent choice history, we fitted multilevel logistic regression models predicting the switch vs. repeat choice by the player’s and opponent’s switch history from *n*-3, *n*-2, and *n*-1 trials, the overall switch probability of the opponent, whether trial *n*-1 was a win or a loss trial, and the interactions between win/loss and all history/context variables (i.e., 15 predictors in total). We estimated both fixed and random effects of all predictors. For Experiment 4, the model had three levels in which trials were nested within players, which in turn were nested in dyads. For the other experiments, models included only the first two levels. The signature of model-based selection is that predictors representing the opponent’s switch rate (e.g., the overall switch probability and opponent’s switch history) is positively related with the player’s switch probability on post-win trials and negatively on post-loss trials. The main prediction we wanted to test was that following post-win trials the predictive relationship is stronger than following post-loss trials. Therefore, we examined the interaction between the post-win/loss contrast and each opponent-related predictor after reversing the label of the predictor for post-loss trials (e.g., *n*–1 opponent “switch” is relabeled as “repeat”, 80% overall switch rate becomes 20%). This allowed us to test the difference in the strength of the relationship, while ignoring the direction of the relationship. We had no a-priori prediction about the direction of the relationship between previous switch/repeat choices and the trial *n* switch/repeat choice. Nevertheless, for a conservative test of post-win/loss differences we again reversed the post-loss label in each case where there was an empirical flip in sign between post-loss and post-win coefficients. We also present in Figure S6 the results without reversing labels.

#### EEG Recordings

In Experiment 5, Electroencephalographic (EEG) activity was recorded from 20 tin electrodes held in place by an elastic cap (Electrocap International) using the International 10/20 system. The 10/20 sites F3, Fz, F4, T3, C3, CZ, C4, T4, P3, PZ, P4, T5, T6, O1, and O2 were used along with five nonstandard sites: OL midway between T5 and O1; OR midway between T6 and O2; PO3 midway between P3 and OL; PO4 midway between P4 and OR; and POz midway between PO3 and PO4. The left-mastoid was used as reference for all recording sites. Data were re-referenced off-line to the average of all scalp electrodes. Electrodes placed ∼1cm to the left and right of the external canthi of each eye recorded horizontal electrooculogram (EOG) to measure horizontal saccades. To detect blinks, vertical EOG was recorded from an electrode placed beneath the left eye and reference to the left mastoid. The EEG and EOG were amplified with an SA Instrumentation amplifier with a bandpass of 0.01–80 Hz and were digitized at 250 Hz in LabView 6.1 running on a PC. We used the Signal Processing and EEGLAB (44) toolboxes for EEG processing in MATLAB. Trials including blinks (> 60 μv, window size = 200 ms, window step = 50 ms), large eye movements (> 1°, window size = 200 ms, window step = 10ms), and blocking of signals (range = −0.01 μv to 0.01 μv, window size = 200 ms) were rejected excluded from further analysis.

#### EEG Analysis

Single trial EEG signals were segmented into 1250 ms epochs starting from 200 ms before the onset of feedback. Thus, each epoch included 700 ms post-feedback periods and initial 250 ms intervals of next trials. After baselining signals with data from initial 200 ms intervals, EEG activity from electrodes Fz and Cz was averaged and transformed into a z-score within subjects. These electrodes were selected based on previous studies reporting a robust interaction between the feedback and the probability context during reinforcement learning (43). Each electrode’s EEG signal was also prewhitened by linear and quadratic trends across trials and blocks. The resulting signal was regressed using a multilevel model with two levels (i.e., electrodes nested within trials) on context variables, as described in the Results section. For illustrative purposes, this was done for every time sample (see Figure 4). In addition, for statistical tests of the post-win versus post-loss regression coefficients, the analyses predicting choices, and the topographic maps, we averaged the EEG signal for an a-priori defined a 300-700 ms interval from the post-feedback period, which was derived from the literature on ERP feedback effects (29, 43). The difference between post-win/loss models was tested in the same manner as in the multilevel model for history effects, namely by reversing predictors of opponents’ history/context to obtain the interaction effect of post-win/loss (see *History Analysis* section).

### Additional Results

#### Rate of Success

Participants’ switching behavior as function of the simulated opponents’ switch rate and *n*-1 win/loss feedback indicates that they used a model of the opponent mainly on post-win, but to a lesser degree on post-loss trials. Obviously, model-based behavior can be useful only when the opponent exhibits some degree of regularity. Therefore, we expect that participants show a greater success rate both after win than after loss feedback and when the opponent’s switch rate deviates from chance (*p*=.5).

**Fig. S1.**
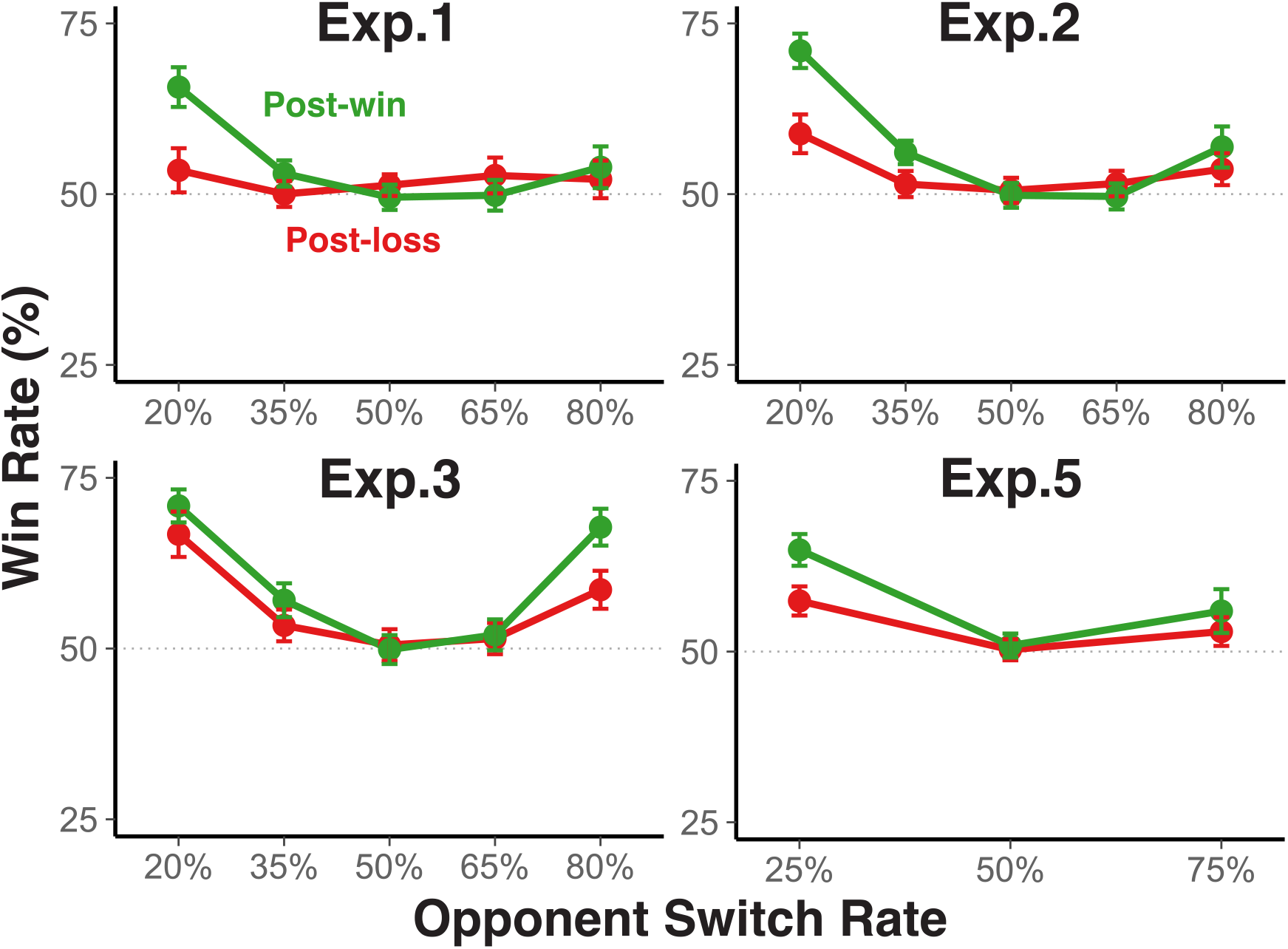
Rate of winning as a function of *n*-1 wins versus losses and opponents’ switch rate. Results for Experiment 2 are collapsed across the two RSI conditions, which showed almost identical results. Error bars show 95% within-subject confidence intervals.

As shown in Figure S1, we found such a pattern for all four experiments with simulated opponents: The success rate followed a right-tilted, U-curved function with the most wins for the lowest switch rate, followed by the highest switch rate, and then the mid-range switch rates. In addition, this pattern was much more robust for post-win than for post-loss trials. The fact that rate of winning was highest for the opponent with lowest switch rate, in particular after win trials is consistent with the fact that participants showed greater tendency for model-based behavior when it required them to engage in low rather than high rates of switching (see Figure 2). Within each experiment, the main effect of *n*-1 wins versus *n*-1 losses was highly significant (all *F*s>= 24.5, *p*<.001), as was the interaction between this factor and the quadratic trend for opponent switch rates (all *F*s>= 10.77, *p*=0.003).

#### Action Choices

Traditionally, when analyzing choice behavior in experimental games, the focus is on the how players choose between different options. Given that our behavioral signature for model-based and stochastic behavior was based on the rate of switching between action choices, we focused on the switch/repeat choice as our primary dependent variable. To ensure that we are not missing important results by only focusing on switch rate, we also examined for all experiments the allocation of choices between the “freeze” and the “run” option (or “up” and “down” for Experiment 3), as well as the degree to which choices were affected by our key independent variables (post-win/post-loss and opponent switch rate). In all experiments, the choices were fairly evenly distributed (i.e., close to 50% for either option). The independent variables had at best only very small effects that were not consistent across experiments. Figure S2 shows example results from Experiment 1; the pattern for the remaining experiments is very similar. Thus, there are no obvious results in the pattern of action choices that would qualify the conclusions form the switch-rate data.

**Fig. S2.**
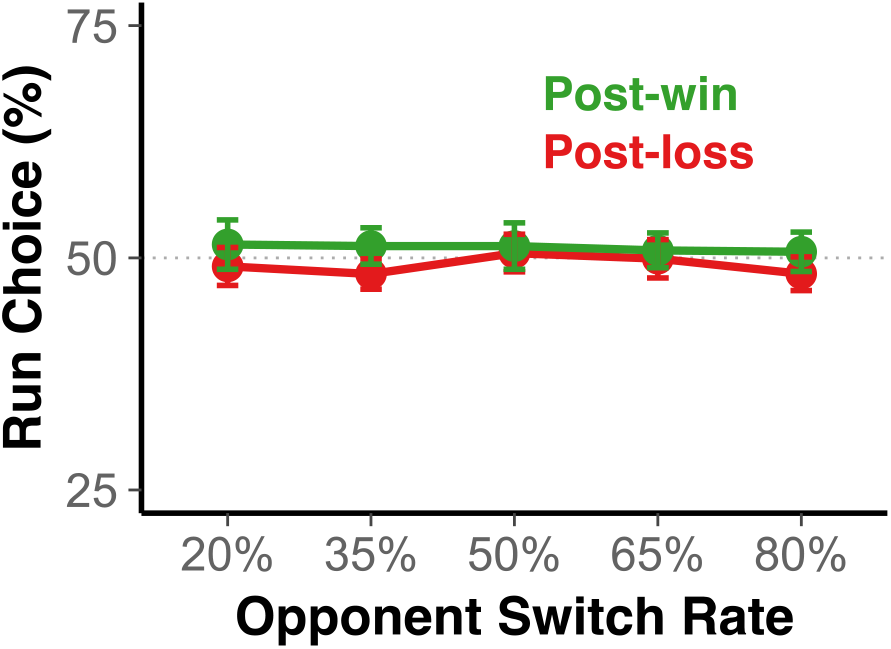
Probability of choosing the “run” option as a function of *n*-1 wins versus losses and opponents’ switch rate for Experiment 1. Error bars show 95% within-subject confidence intervals.

#### Are Feedback Effects Temporary?

Our model assumes that the effect of loss-feedback does not eliminate the model of the opponent, but rather depresses it temporarily. Thus, we should expect that that win-loss feedback has a large effect on the next-trial choice, and either no, or only a small effect thereafter. Figure S3 shows for Experiment 1 the switch-rate function from Figure 2, but further conditioned on the trial *n*-2 win-loss feedback. As apparent, choice behavior is dominated by the effect of trial *n*-1 feedback. There was is a small additional effect of trial *n*-2 feedback, such that model-based behavior is strengthened following two consecutive wins and stochastic behavior is strengthened following two loss trials (i.e., after two win-trial in a row, the switch-rate function slope become more positive, after two loss-trials the function becomes more shallow). Analyzing these data with an ANOVA with the factors trial *n*-2 and trial *n*-1 feedback as well as a linear contrast for the opponent switch-rate factor, revealed a strong n-1 feedback x switch-rate interaction, *F*(1,51)=58.45, *p*<.001, *eta^2^*=.53, and a much weaker, but still reliable *n*-2 feedback x switch-rate interaction, *F*(1,51)=15.02, *p*<.001, *eta^2^*=.23, and no three-way interaction, *F*(1,51)=.25, *p*=.91. The results from the remaining experiments were similar to this pattern, but generally showed slightly weaker n-2 feedback effects. The fact that there was a small, cumulative effect of trial *n*-2 feedback indicates some degree of adaptation to consistent win or loss feedback contingencies. Yet, the fact that choices are mainly dominated by trial *n*-1 feedback indicates that loss feedback only temporarily depresses the model representation, allowing quick recovery of the last-used model representation following a subsequent win.

**Fig. S3.**
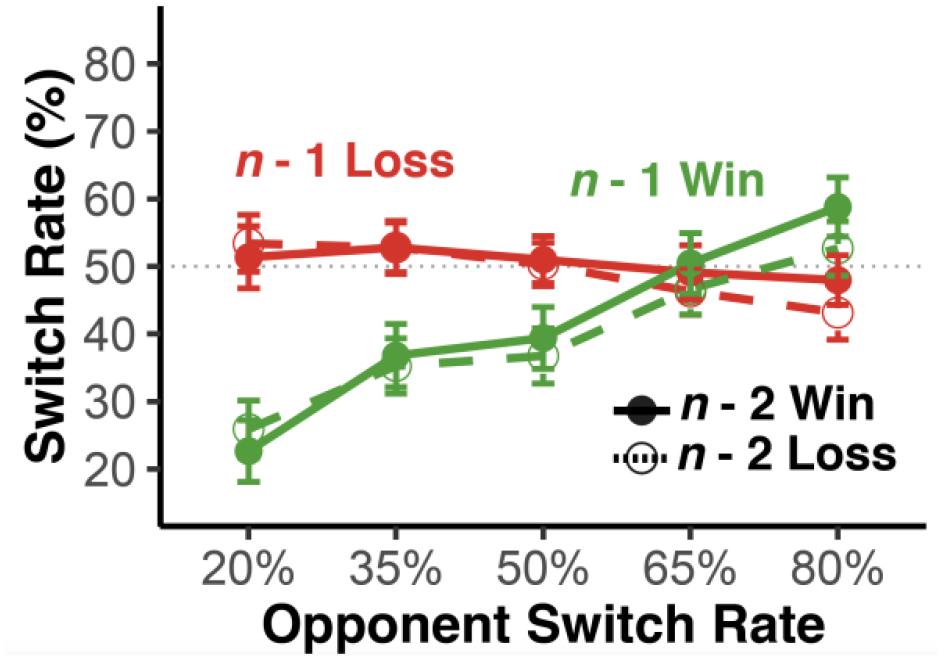
Empirical switch rate as a function of opponents’ switch rate and both trial *n*-1 win versus loss feedback and trial *n*-2 win versus loss feedback for Experiment 1. Error bars show 95% within-subject confidence intervals.

#### History Analyses

Our analyses to test the win/loss difference in the strength of the relationships between history/context and current choices involved selectively reversing labels of some of the predictors (see Figure 3). In Figure S4, we show the, original, signed coefficients. As apparent, for the opponent-related predictors, coefficients were generally positive following win feedback and negative, albeit smaller in size, following loss feedback. This pattern is consistent with our mixed-strategy account, by which choice is model-driven after win trials, but less influenced by the opponent’s global or local choice context following losses.

Our predictions were mainly with regard to the influence of opponents’ choices and we had no clear expectations about how the player’s own choice history would influence future choices. In fact, the pattern of player-related effects was not as consistent as for the opponent-related predictors. However, with few exceptions, the strength of the effects seemed stronger following win than following loss trials. This result is consistent with the conclusion that loss-feedback dampens the influence of the recent task context in general, not just as it relates to the opponents’ behavior.

We also conducted a more straightforward assessment of the effect of history on switch/repeat choices that did not require selective recoding of predictors. Specifically, we performed individual, logistic regression analyses for each subject, and separately for post-win and post-loss trials. Given that the effect of overall switch context was already demonstrated in our initial analyses (see Figure 2), we only included here the players’ and the opponents’ trial *n*-1 to *n*-3 switch/repeat choices as predictors. Figure S5 shows for each experiment the post-win and post-loss distributions of *R*^2^ scores resulting from these analyses (45). As apparent, post-win distributions were in all cases significantly farther to the right than post-loss distributions. Thus, these analyses confirm that following wins, switch/no-switch decisions are overall more dependent on history than post-loss decisions.

**Fig. S4.**
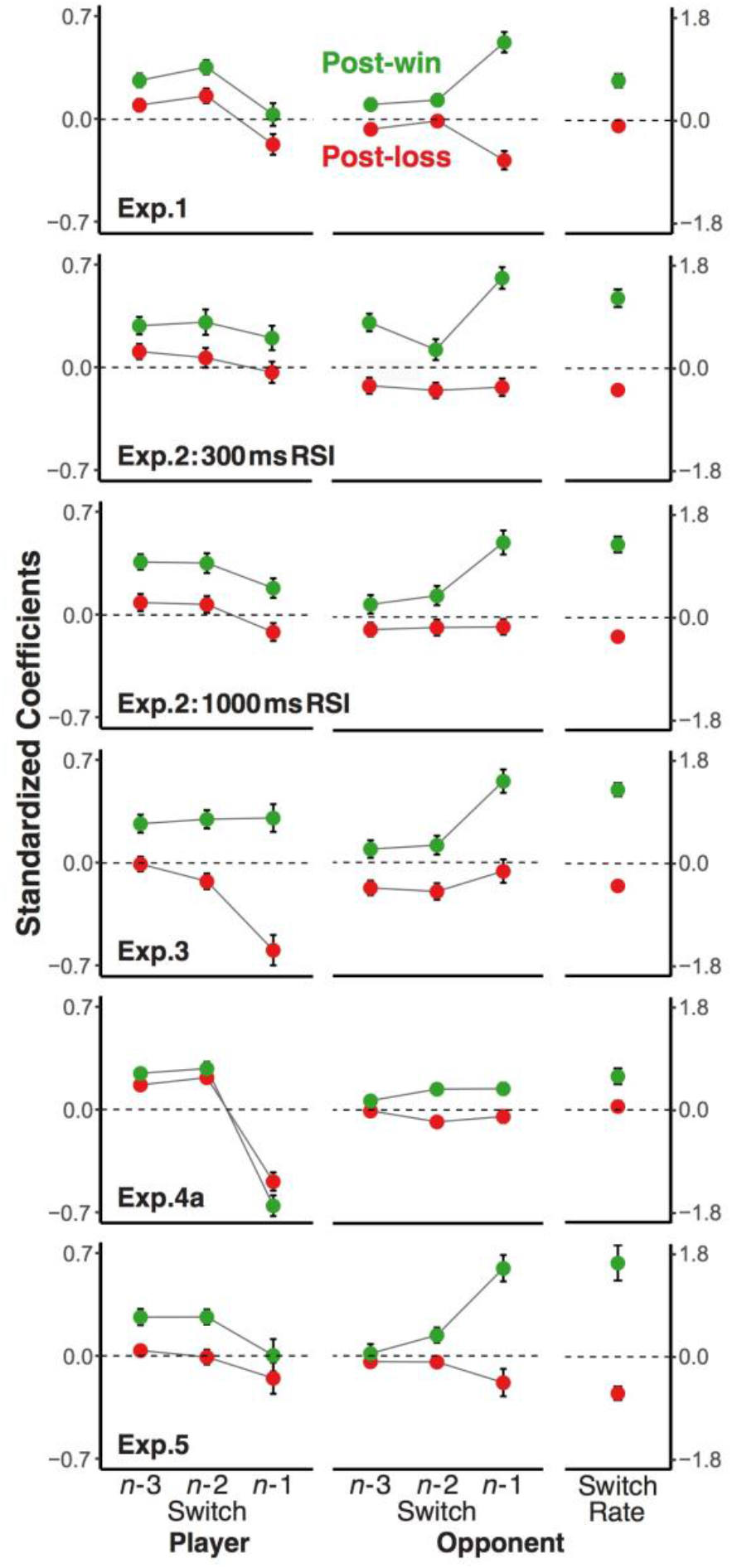
Signed standardized coefficients from multi-level logistic regression models predicting the trial *n* switch/repeat choice on the basis of players’ and opponents’ switch/repeat choices on trials *n*-1 to *n*-3 and the opponents’ overall switch rate, run separately for post-win and post-loss trials. Error bars are within-subject standard errors around the coefficients.

**Fig. S5.**
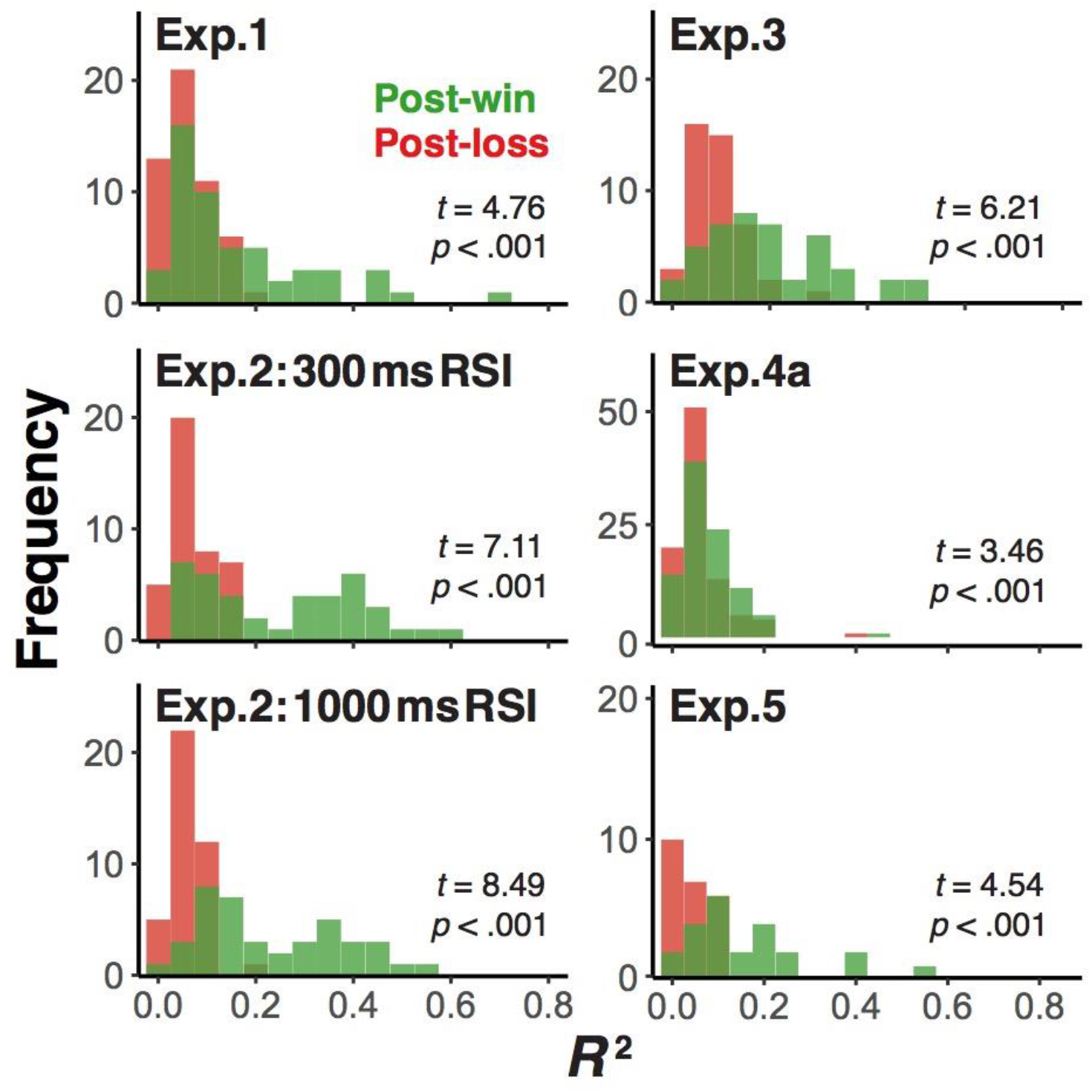
Histogram of Cox and Snell pseudo *R*^2^-square scores from logistic regression models fitted within subjects (dark green shading indicates overlapping regions of the distributions). The difference in fit scores between post-win/loss was tested via t-test after converting *R*^2^ values into *z*-score. Fit scores are generally higher for post-win models compared to post-loss models.

#### Modelling of Switch Choices

Figure 2 in the main paper contains predictions from the following probabilistic model that captures four different, potential influences on subjects’ average switch rates *p*_switch_ across conditions:

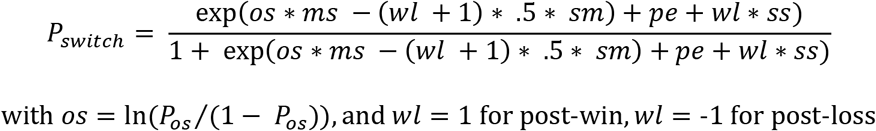

where *p_os_* is the experimentally manipulated probability of a switch by the simulated opponent, which is translated into its log-odds form (*os*), and *wl* codes for wins versus losses on trial *n*-1. The parameter *ms* (***m****odel **s**trength*), represents the strength of the internal model (e.g., *m*s=1 would indicate direct probability matching between the opponent’s and the player’s switch probability). The parameter *sm* (***s****uppression of **m**odel*), within the term (*wl*+1) * .5 * *sm* represents the degree to which the model-based choice is changed on post-loss relative post-win trials. A negative *sm* parameter would indicate suppression of the model in favor of stochastic choice following losses. In addition, a positive *pe* (***p****erseveration **e**ffect*), parameter would represent the tendency to unconditionally favor the previously choses task; and a positive *ss* (*win-**s**tay/lose-shift*), parameter indicates that choices are dominated by the win-stay/lose-shift strategy.

We applied this model both to the average data for each experiment, and to the subject-specific averages. Table S1 shows the estimated parameters for each of the four experiments, their confidence intervals, as well as model fits (*R*^2^) for the group-average data and also the parameters and confidence intervals for the averages of the individual-specific estimates. Across experiments and individuals, each of the four different influences are relevant for characterizing participants’ propensity to switch from one trial to the next. For group-average data, the model strength parameter ranged between .48 and .87, indicating that overall, the opponent’s switch rate affected the participant’s switch rate in an incentive-compatible manner. Average *ms* values below 1.0 also indicate that participants overall engaged in “imperfect” probability matching (a value of 1.0 would indicate perfect probability matching, values above 1.0 a maximizing tendency). There is a substantial literature indicating that probability matching is the dominant, albeit suboptimal strategy in serial decision tasks (46, 47).

**Table S1.** Parameter estimates and 95% confidence intervals from fitting the choice model to group average and individual data from Experiments 1, 2, 3, and 5.

|  | Fitting Group Averages |  |  |  |  | Fitting Individuals' Data |  |  |  |
| --- | --- | --- | --- | --- | --- | --- | --- | --- | --- |
| Parameters | *ms* | *sm* | *pe* | *ss* | $R^2$ | *ms* | *sm* | *pe* | *ss* |
| *Simulated Opp.* |  |  |  |  |  |  |  |  |  |
| Exp. 1 | .48±.10 | -.38±.09 | .21±.06 | .20±.06 | .975 | .61±.19 | -.50±.16 | .24±.14 | .22±.11 |
| Exp. 2, 300 ms | .74±.12 | -.47±.15 | .28±.07 | .30±.07 | .988 | .96±.26 | -.66±.24 | .34±.10 | .36±.14 |
| Exp. 2, 1000 ms | .74±.13 | -.49±.17 | .19±.08 | .26±.08 | .984 | .93±.24 | -.67±.21 | .25±.11 | .33±.13 |
| Exp. 3 | .86 ±.11 | -.44±.14 | .07±.04 | .24±.06 | .993 | 1.13±.28 | -.68±.23 | .11±.10 | .29±.11 |
| Exp. 5 | .87±.17 | -.50±.21 | .11±.09 | .30±.09 | .998 | 1.10±.39 | -.71±.30 | .16±.10 | .36±.24 |
| *Human Dyads* |  |  |  |  |  |  |  |  |  |
| Exp. 4 |  |  |  |  |  | .16±.10 | -.14±.13 | .11±.9 | .31±.9 |

*Note.* *ms*=model strength, *sm*=suppression of model, *pe* = perseveration effect, *ss* = win-stay/lose-shift tendency. For Experiment 2, fits are reported separately for the 300 ms and the 1000 ms RSI condition. Fits for individual subjects in Experiments 1, 2, 3, and 5 are on the basis of each subject's condition averages. For Experiment 5, we report parameters resulting from modeling individuals' trial-by-trial choices.

The lower-level perseveration and win-stay/lose-shift biases had modest, but statistically robust effects on choice. For example, the *pe* of .21 found in Experiment 1 translates into a switch probability of *p*=.45, when all other factors are ignored––a small overall perseveration bias. Similarly, the *ss* of .20 found in Experiment 1 implies a switch probability of *p*=.45 following wins and of *p*=.55 following losses, again all other factor ignored. Most importantly, the theoretically critical, strategy mixture parameter *sm* ranged from -.38 to -.50 and was reliably lower than 0 across all experiments. These parameters estimates implied that on 50-80% of post-loss trials, the model-based influence was replaced by stochastic choice.

The individual-specific parameter estimates also allowed us to examine the degree to which the different influences on choice were tied to successful competitive behavior. To this end, we entered each individual’s four different parameter estimates as fixed-effect predictors into a two-level regression analysis with experiment as random factor and overall success as criterion variable. While on average, *pe* and *ws* indicated the expected perseveration and win-stay/lose-shift biases (i.e., *pe*<0 and *wl*<0), there were substantial individual differences in these parameters that included individuals with alternation or win-shift/lose-stay biases (i.e., *pe*>0 and *ss*>0). Given that any bias can imply a deviation from optimal performance, we coded these two parameters in absolute terms.

**Table S2.** Using parameter estimates from the choice-model fitted to individual’s condition means to predict individual’s competitive success.

|  | *b* | *se* | *t*-value |
| --- | --- | --- | --- |
| intercept | .504 |  |  |
| *ms* | .086 | .007 | 12.31 |
| *sm* | .064 | .008 | 8.03 |
| abs(*pe*) | -.016 | .006 | -2.56 |
| abs(*ss*) | -.001 | .001 | -0.10 |

*Note.* Shown are raw, fixed-effect coefficients (*b*), the standard error around the coefficients (*se*), and the associated *t*-value. Experiment was coded as a random grouping factor. Absolute values for the *pe* and the *wl* effect were used to account for biases in either direction. Note, that the more negative the *sm* parameter, the greater the suppression of model-based choice on post-loss trials. Thus, a positive coefficient in this analysis indicates that more suppression leads to worse performance.

As shown in Table S2, model strength has a highly robust positive effect on success, whereas either a perseveration or an alternation bias reduced the amount of money earned; no corresponding effect was found for the win-stay/lose-shift parameter. As would be expected, the main effect of the strategy-choice parameter was positive, implying that less stochastic behavior after losses produced greater overall success.

#### Modeling of Switch Choices in Experiment 4a (Dyads)

In Experiment 4a, each participant played against another participant within seven, 80-trial blocks. Different from the other experiments, where the simulated opponent’s behavior was under experimental control, we here had to utilize the natural, within-session variability of each player’s switch rate within a trial-by-trial version of our choice model. For this purpose, the *p*_os_ parameter, which represents the opponent’s choice behavior was calculated as a running average of the opponent’s switch rate within each block. The ending running average of block *n*-1 (or *p*=.5 for block *n*=1) was used as a starting value for block *n*, allowing some carry-over of prior knowledge of the opponent’s previous-block behavior.

Results from this model are shown in the bottom row of Table S1. Not surprisingly, the model strength parameter *ms* was substantially smaller than in the preceding experiments, but still significantly larger than 0. Also, the perseveration parameter *pe* and the win-stay/lose-shift parameter *ss* were robust and roughly in a similar range as in the remaining experiments. Importantly, the theoretically critical suppression of model parameter *sm*, was also statistically significant and of about the same size as the model-strength parameter *ms*, indicating that on post-loss trials the effect of the model is essentially eliminated.

We also used a multi-level regression model with participants grouped within dyads to predict each participant’s success (in terms of probability of win trials) as a function of the four model parameters. As in the preceding model (Table S2), we again used absolute values from the perseveration and the win-stay/lose-shift scores in order to capture biases in in either direction (very similar results would have been obtained with signed values). As shown in Table S3, greater reliance on the model, a smaller tendency to disregard the model after losses (i.e., a less negative *sm* score), a smaller, absolute perseveration score, and a larger absolute win-stay/lose-shift all contributed to greater success. Aside from the result for the win-stay/lose-shift score, the overall pattern was qualitatively very similar to the results from the simulated-opponent experiments. The relatively strong positive link between the win-stay/lose-shift bias and success rate is noteworthy. Exploring it further goes beyond the scope of this paper. However, it is important to emphasize that these analyses do not reveal the causal pathway and therefore do not necessarily imply that a stronger win-stay/lose-shift bias leads to greater success. It is just as plausible that players who are more successful are more inclined to repeat successful moves, which then might go unnoticed by their weaker opponents.

**Table S3.**
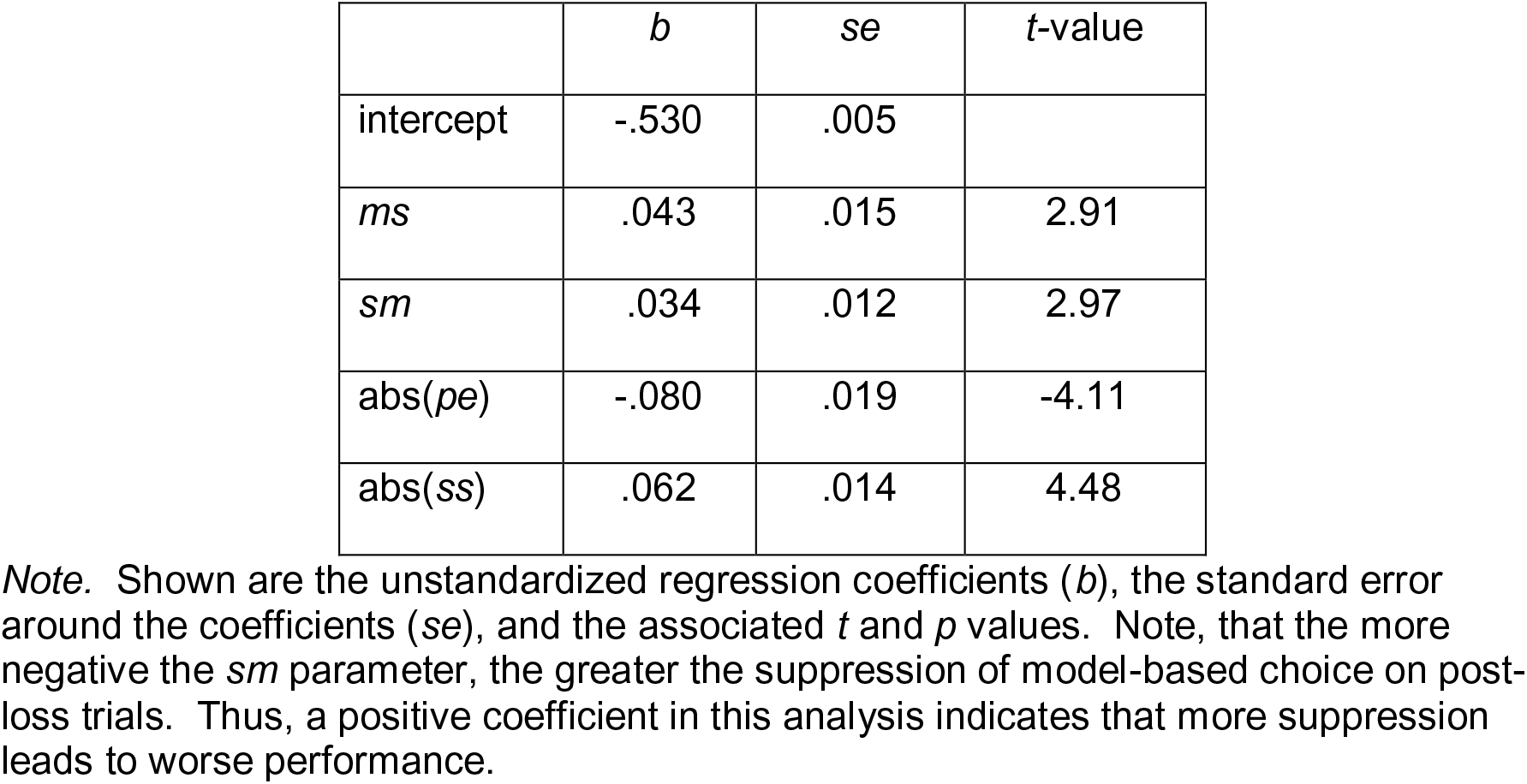
Using parameter estimates from the choice-model fitted to individual’s trial-by-trial data to predict the proportion of win trials (n=94) in Experiment 4a.

#### Event-related Potentials

Our key interest in Experiment 5 was in the amount of information about history/context variables that was contained in the EEG signal following the feedback signal, and how that information predicted behavior. Therefore, we present in Figure 4 analyses that capture the degree to which history/context variables are expressed in the EEG signal, using coefficients from multi-level regression analyses. These analyses are adequate as we had no strong a-priori predictions about how the different context variables and interactions between them would be expressed in the EEG signal and the regression-based analyses yielded trial-by-trial estimates of “representational strength” that we used in the PPI analyses (see below). Nevertheless, it can be informative to examine how the different variables affect the EEG signal when examined in the traditional manner, namely in terms of event-related potentials (ERPs).

Figure S6 shows the ERPs from all relevant conditions. Following the feedback (200 to 300 ms after the onset), we observed a typical feedback-related negativity (FRN), with a peak that was more negative for loss feedback compared to win feedback. Consistent with the results of multi-level modeling (Figure 4 and Figure S6), the amplitude of the FRN was affected by the combination of feedback and context variables. Specifically, it appears that the FRN amplitude was most negative for unexpected opponent switch or repeat choices, that is for opponent switch choices for 25% switch-rate opponents and (to a lesser degree) for repeat choices when facing 75% switch-rate opponents). These effects in turn, were considerably more pronounced following win than loss trials.

**Fig. S6.**
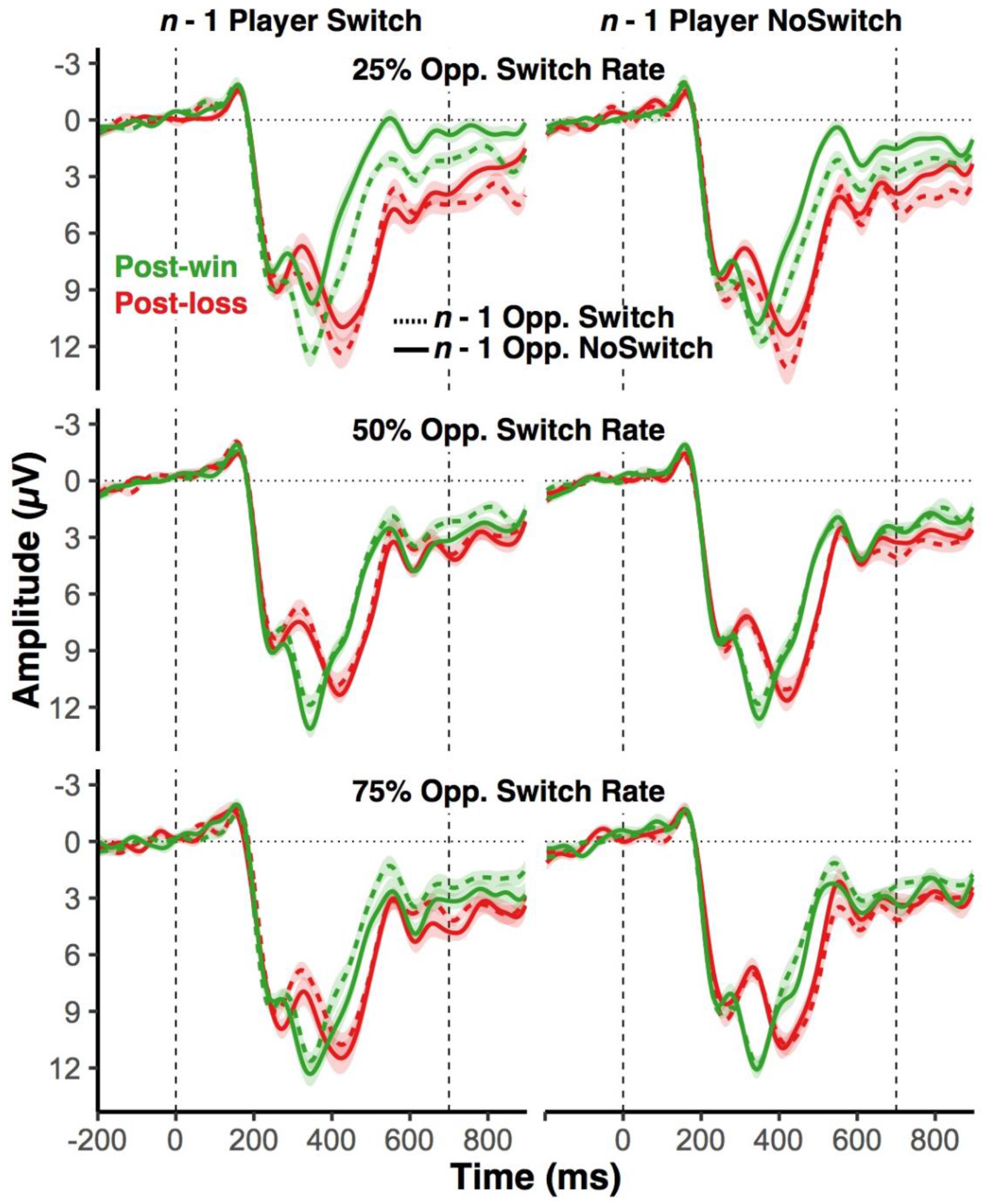
Event related potentials (grand average EEG activity for electrodes Fz and Cz) grouped by all factors used in the EEG analysis: the opponent’s overall switch rate (20,50,75%), the *n*-1 opponent switch/no-switch choice, and the *n*-1 player’s switch/no-switch choice. The EEG signal was low-pass filtered (Butterworth, 25*Hz*), time-locked to the onset of feedback, and subtracted from the average across the 200 ms baseline period prior to the feedback signal. The shaded area indicates within-subject 95% confidence intervals around the average signal.

Within the large literature on feedback-related EEG effects (31, 48–50), there is no general agreement to what degree the main components (in particular the FRN) reflect an unsigned prediction error (i.e., surprise) or a signed prediction error (i.e., a reinforcement signal). The fact that we generally find a larger negativity during the typical FRN time range (200 to 300 ms after the onset of feedback) following loss trials is consistent with a negative prediction error account. However, the fact that we also see a clear expectancy modulation of the FRN for post-win trials (i.e., a larger positivity when opponents with low switch rates do switch or opponents with high switch rate do not) is also consistent with the FRN as an unsigned prediction error. The literature does not provide much information on the degree to which context modulates the FRN signal differentially for negative versus positive feedback (32). One interesting exception is a study by Cohen et al. (10) who reported that learned reward expectations decreased the FRN amplitude selectively on gain trials, but not on loss trials. Thus, as in our results, this finding suggests that expectations about task contingencies are expressed more strongly after positive than after negative feedback in the feedback-related EEG signal.

#### Controlling for Upcoming Switch

Figure 4 shows the degree to which history/context variables are expressed in the EEG. It is possible that the feedback-related differences for the history/context coefficients shown in Figure 4 are due to the fact that feedback affects the probability of an upcoming switch. Therefore, the EEG effects may reflect preparatory processes associated with an upcoming switch, such as the allocation of effort. We show in Figure S7 results from the same analyses presented in Figure 4, but in addition entering the switch on the next trial (coded 0/1) as predictor. As evident, here was indeed a small, significant relationship between the EEG signal and the upcoming switch. However, the overall pattern of history/context effects remains the same even when controlling or this variable.

**Fig. S7.**
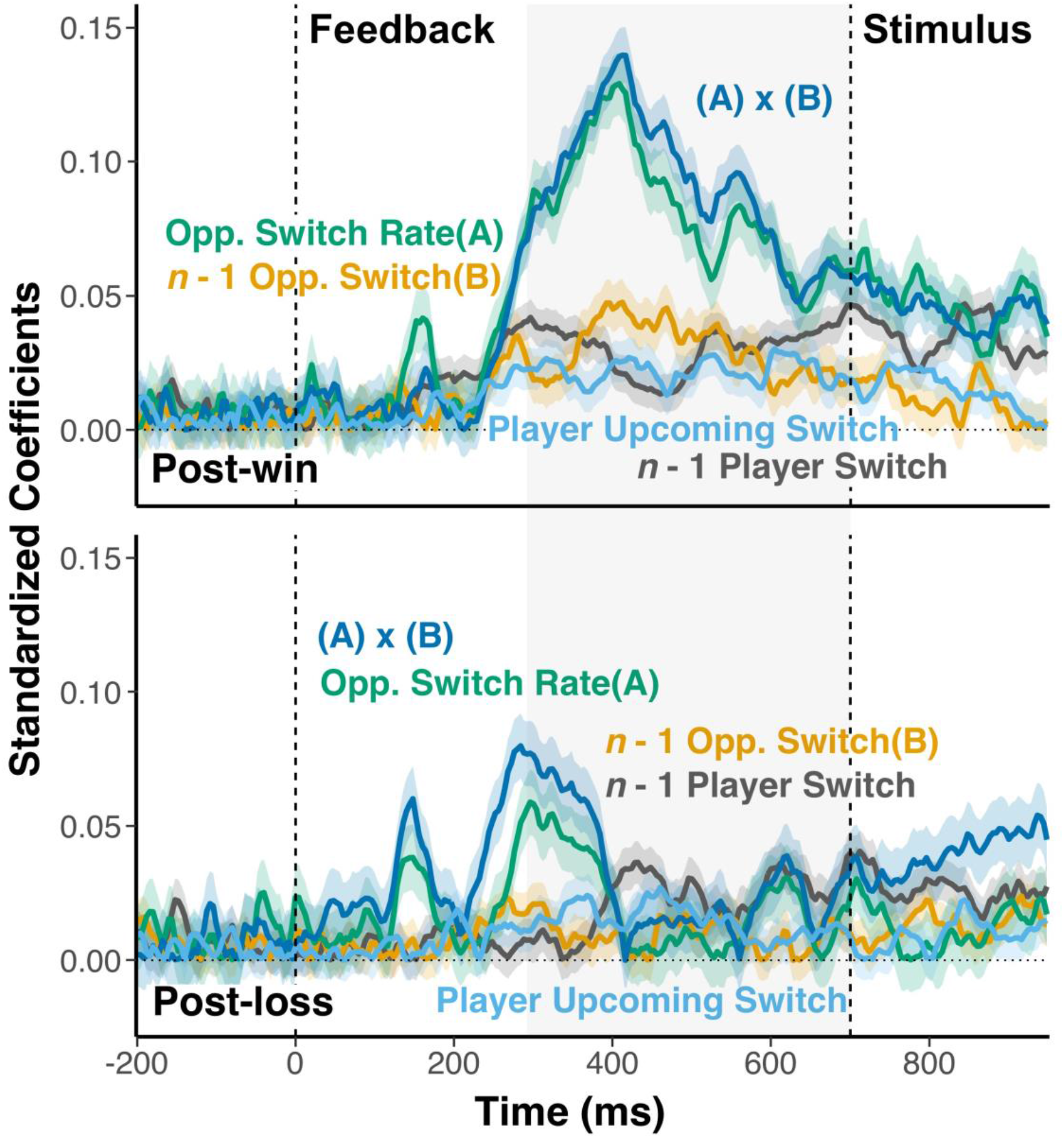
Standardized coefficients from multi-level regression models relating EEG activity at Fz and Cz electrodes to main predictors and the control variable (players’ upcoming switch). Shaded areas around each line indicate within-subject standard errors around coefficients.

**Fig. S8.**
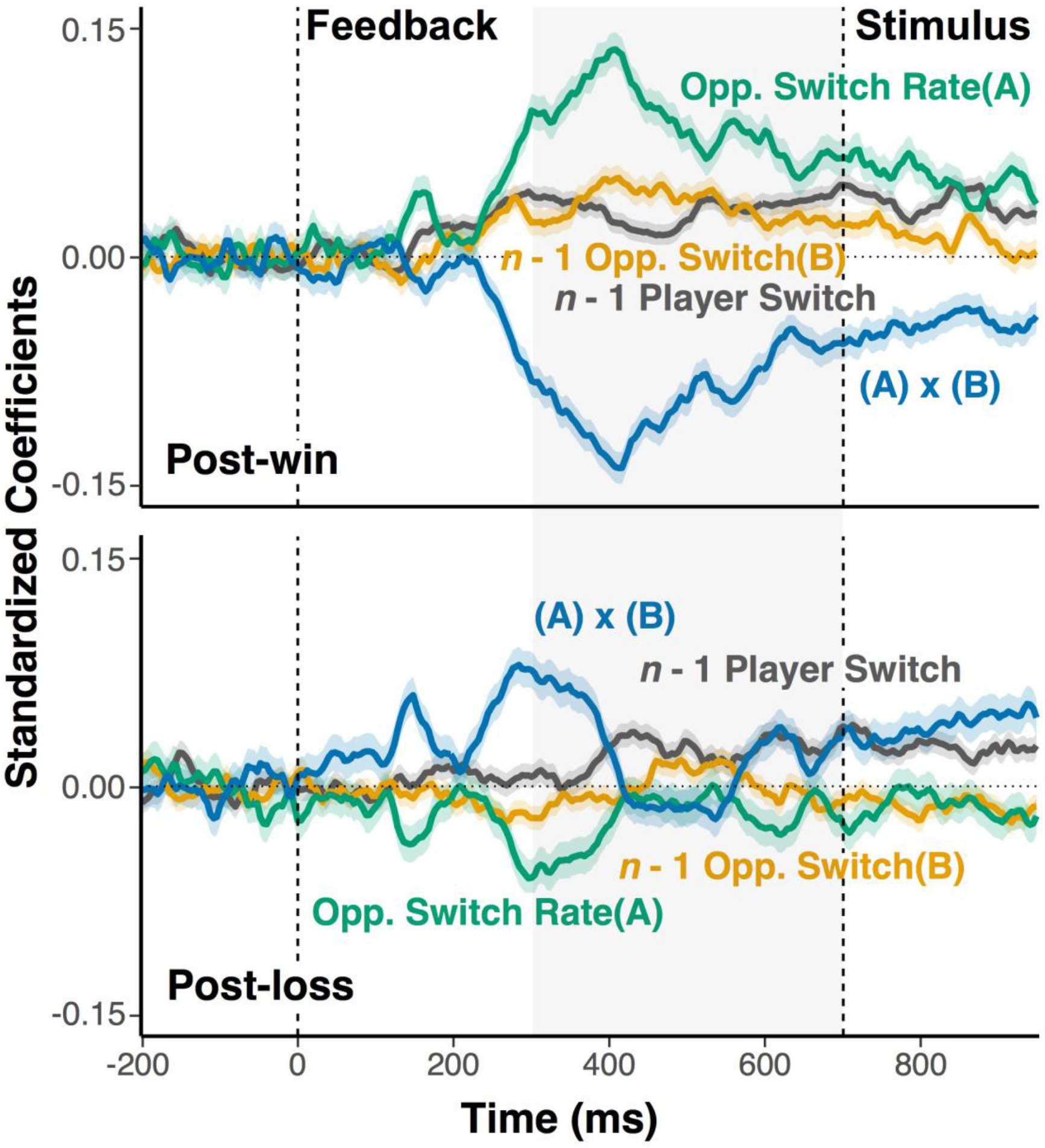
Signed standardized coefficients from multi-level regression models relating EEG activity at Fz and Cz electrodes to the opponent’s overall switch rate (A), the *n*-1 opponent switch/no-switch choice (B), the *n*-1 player’s switch/no-switch choice, and the interaction between A) and B) for each time point and separately for post-win and post-loss trials. Shaded areas around each line indicate within-subject standard errors around coefficients.

#### Signed Coefficients

In Figure 4, we had reversed the labels for the opponent-related predictors because our predictions referred to the amount of information about the competitive context, not how exactly that information is expressed in the EEG signal. Figure S8 presents the identical analyses as in Figure 4, however plotted using the original predictors (i.e., without reversing labels on post-loss trials). Inspection of the signed coefficients reveals a clear distinction between post-loss and post-win signals. For all opponent-related predictors, the effect on the EEG signal was not only reduced following losses, it was also flipped in sign relative to post-win trials. Note, that opponents’ local and global switch behavior has very different implications for the subject’s behavior depending on whether one is currently winning or losing (e.g., see Figure 1b). Thus, one might speculate that this flip in sign is indicative of the win/loss-contingent difference in interpretation (or behavioral implication) of the information provided through the opponent.

#### PPI Analysis

In the main article, we present a psychophysiological interaction (PPI) analysis that predicts trial-by-trial switch choices from the degree to which history/context is expressed in the EEG signal. Given that we only found a robust representation of context in the EEG signal following win trials (Figure 4), we restricted our analysis to these trials. Note also that the theoretical status of post-loss trials is much less obvious than that of post-win trials. Given the reduced context representation following losses, there may simply be too little meaningful variability to establish a reliable relationship with behavior. At the same time, the remaining variability might also still have a detectable influence, just with a smaller overall effect on behavior. Therefore, there is no strong, theoretical reason to test post-win and post-loss relationships against each other. Nevertheless, in Table S4 we show the results of PPI analyses separately for post-win and post-loss trials. The results indicate that the relationship between the EEG representation of the global, opponent switch rate and behavior was equally strong for post-win as for post-loss trials. Note, that the combination between the identical sign for the EEG-behavior relationship across post-win and post-loss trials and the flipped sign for the opponent switch rate/EEG relationship (Figure S8) is consistent with the reversal of relationship between opponent, overall switch rate and player switch rate depending on win or loss feedback (e.g., Figure 2). For the local, history variables, we found robust effects only following win trials, but not following loss trials.

**Table S4.** Coefficients from the PPI analysis predicting upcoming choices using residuals of MLM regression model for post-win and post-loss trials.

|  | Post-win |  |  | Post-loss |  |  |
| --- | --- | --- | --- | --- | --- | --- |
|  | *b* | *se* | z-value | *b* | *se* | z-value |
| residual EEG | 0.20 | 0.03 | 6.39 | 0.07 | 0.04 | 2.05 |
| Opponent Switch Rate(A) | 0.11 | 0.05 | 2.26 | -0.11 | 0.05 | -2.37 |
| *n*-1 Opponent Switch (B) | -0.13 | 0.04 | -3.36 | -0.03 | 0.03 | -0.95 |
| *n*-1 Player Switch | -0.11 | 0.03 | -3.36 | -0.12 | 0.03 | -0.36 |
| (A) x (B) | -0.06 | 0.06 | -0.89 | 0.08 | 0.06 | 1.37 |

*Note.* Shown are the unstandardized regression coefficients (*b*), the standard error around the coefficients (*se*), and the associated z values.

